# Effects of cell growth and temperature on mitochondrial physiology of *Drosophila melanogaster* embryonic cell line

**DOI:** 10.1101/2025.11.25.690158

**Authors:** Rodiésley S. Rosa, Ana Paula M. Mendonça, Matheus P. Oliveira, Danielle B. Carvalho, Leonardo Vazquez, Marcus F. Oliveira

## Abstract

The fruit fly *Drosophila melanogaster* is a valuable model for studying cellular and metabolic processes conserved with mammals. Using its embryonic Schneider S2 cell line, commonly used for heterologous protein expression, we investigated how culture duration and slight temperature changes affect mitochondrial physiology via high-resolution respirometry. We assessed five mitochondrial bioenergetic parameters and two coupling parameters at early and late growth phases, and at two temperatures (26°C and 28°C). We observed that a moderate temperature increase coupled with late growth phase synergistically enhanced basal, OXPHOS, and leak respiration, with only minor impacts on mitochondrial coupling efficiency. Notably, spare and maximal respiratory capacities increased disproportionately with either cell growth progression or moderate temperature rise, suggestive of enhanced mitochondrial biogenesis. These results demonstrate that small temperature increases and cell maturation activate mitochondrial bioenergetic capacity through mechanisms consistent with mitochondrial biogenesis and functional remodeling. Such adaptations improve cellular metabolic resilience while maintaining energy conversion efficiency. These findings deepen understanding of mitochondrial plasticity in response to environmental and developmental signals. They may inform strategies to optimize mitochondrial physiology in various biological contexts, including aging, metabolic diseases, and stress adaptation.

## 1. Introduction

*Drosophila melanogaster* is a powerful and well-established model organism for understanding fundamental biological processes, including many conserved metabolic pathways relevant to human health (Pandey and Nichols, 2011). The fruit fly possesses organs, such as the insect fat body, that are metabolically and functionally analogous to mammalian tissues like the liver and adipose tissue, serving as key sites for energy storage and endocrine signaling (Gutierrez et al., 2007). This high degree of conservation makes *Drosophila* an invaluable tool for studying complex conditions, including metabolic disorders, cardiovascular disease, and neurodegeneration.

To facilitate cellular-level investigations, cell lines from *Drosophila* embryos were established (Schneider, 1972). Among these, the Schneider-2 (S2) cell line has become a widely used platform. S2 cells, which can grow in suspension or as an adherent monolayer, are extensively employed for studying diverse cellular processes and as a robust system for heterologous protein expression (Abrams et al., 1992; Moraes et al., 2012). These cells enable efficient gene silencing, live-cell imaging of mitosis, and large-scale protein production, making them a valuable tool for functional genomics and host-pathogen interaction research (Cherbas et al., 2011; Elwell and Engel, 2005; Schneider, 1972). Despite their broad application, the fundamental bioenergetics of S2 cells—particularly mitochondrial physiology and its regulation—remain largely unknown (Abrams et al., 1992; Moraes et al., 2012; Pamboukian et al., 2008).

Cellular energy metabolism is dynamically regulated by intrinsic physiological states and external environmental cues. During cell growth and progression through the cell cycle, anabolic demands strongly influence metabolic activity, while factors such as pH and temperature strongly modulate these processes (Humez et al., 2004; Jandova et al., 2012; Place et al., 2017; Vander Heiden et al., 2001). Recent evidence shows that mitochondria can maintain temperatures substantially higher than the surrounding cytosol due to heat generated by OXPHOS (Chrétien et al., 2018), and that mitochondrial enzymes are partially adapted to operate under these elevated thermal conditions.

Several studies have explored how S2 cells respond to temperature fluctuations and stress (Bouchet et al., 2014; Jevtov et al., 2015; Klepsatel et al., 2019; Moraes et al., 2012; Nadeau and Teets, 2020). While one report found no significant difference in maximum specific growth rates between 24°C and 28°C, suggesting stable proliferation within this range (Moraes et al., 2012), other studies revealed activation of stress and metabolic pathways under thermal stress, including the TORC2-mediated heat shock response (Jevtov et al., 2015). Even minor thermal shifts (for example, from 22°C to 24°C) have been shown to cause physiological changes such as alterations in cell size (Jalal et al., 2015).

Consistent with these findings, pharmacological inhibition of OXPHOS in S2 cells maintained at 25°C markedly reduced mitochondrial temperature, as detected by MTY-based ratiometric fluorescence probes (Terzioglu et al., 2023). Altogether, these observations indicate that stable proliferation rates do not necessarily reflect a steady bioenergetic state; instead, cells may balance replication and growth through dynamic adjustments in energy allocation that support different functional requirements.

We hypothesized that minor, physiologically relevant temperature changes (26°C vs. 28°C) and progression through different growth phases would independently and interactively modulate S2 cell mitochondrial physiology. Therefore, the main objective of this work was to investigate the distinct effects of cell growth and minor temperature shifts on the mitochondrial bioenergetic parameters of the S2 cell line.

## 2. Material and methods

### 2.1. Reagents

All general reagents were purchased from Merck-Sigma Aldrich (USA). Fetal Bovine Serum (FBS) and Schneider Medium were obtained from LGC Biotecnologia (Brazil).

### 2.2. Cell culture

Schneider’s *Drosophila melanogaster* embryonic cell line S2 was kindly provided by Prof. Helena Araújo from the Department of Histology and Embryology at the Institute of Biomedical Sciences at Federal University of Rio de Janeiro (UFRJ, Brazil). Cells were maintained in 25 cm^2^ culture flasks at either 26°C or 28°C in Schneider medium supplemented with 10 % FBS. Cells were sub-cultured at 1:34 with 5 mL of media every seven days.

### 2.3. Cell growth

Cells were cultured for experiments in 6-well plates by seeding 8.8 x 10^3^ cells/well, and cell density was determined every two days by removing cells growing in suspension from all wells. Only attached cells were counted and used throughout this study. This standardization is especially important since cells in suspension may exhibit physiological profiles that differ from those of adherent cells (Jang et al., 2022). The adherent cells were then removed by gentle washing with medium and transferred to 15 mL tubes. Cell density and viability were determined over 9 days of culture by taking a small aliquot (∼2 μL) of cell suspension and 1:10 dilution with a 4 % trypan blue and counting in the Neubauer chamber. Overall, Trypan blue staining indicated that less than 5% of the cells were nonviable throughout the 9-day culture period. As a result of the experiments, maximum specific growth rates (*μ_max_*), defined as the highest rate at which cells increase in number per unit of existing cell mass under optimal nutrient and environmental conditions, and maximum cell concentrations of cultures were determined at both temperatures. *μ_max_* values were determined by using the Hill slope sigmoidal dose-response best fit of cell growth curves on GraphPad Prism software version 8.0 for Windows (GraphPad Software, USA).

### 2.4. Respirometry analyses

A volume of media corresponding to 2 x 10^6^ cells was transferred to the chamber of Oxygraph-O2k (Oroboros Instruments, Innsbruck, Austria) filled with 2 mL of Schneider medium without FBS. The cells were allowed to equilibrate for about 10 minutes, with continuous stirring set up at 750 rpm and 27.5°C. Data was generated from at least 3 biological replicates for each study after calibration of the oxygen sensors and instrument background corrections. Oxygen consumption rates were determined by using the DatLab 6.0 software and were normalized by cell numbers. Respiratory rates were registered under routine conditions after signal stabilization and upon different OXPHOS modulators as follows: 250 µmol/ml oligomycin (ATP synthase inhibitor), 300 nM carbonyl cyanide p-(trifluoromethoxy) phenylhydrazone (FCCP) as final concentration, 100 nM rotenone, and 0.5 µg/mL antimycin A. The different mitochondrial metabolic states were then determined by calculating the oxygen consumption rates under the effect of specific OXPHOS modulators (Doerrier et al., 2018), as depicted in Figure 1:

**Figure 1:**
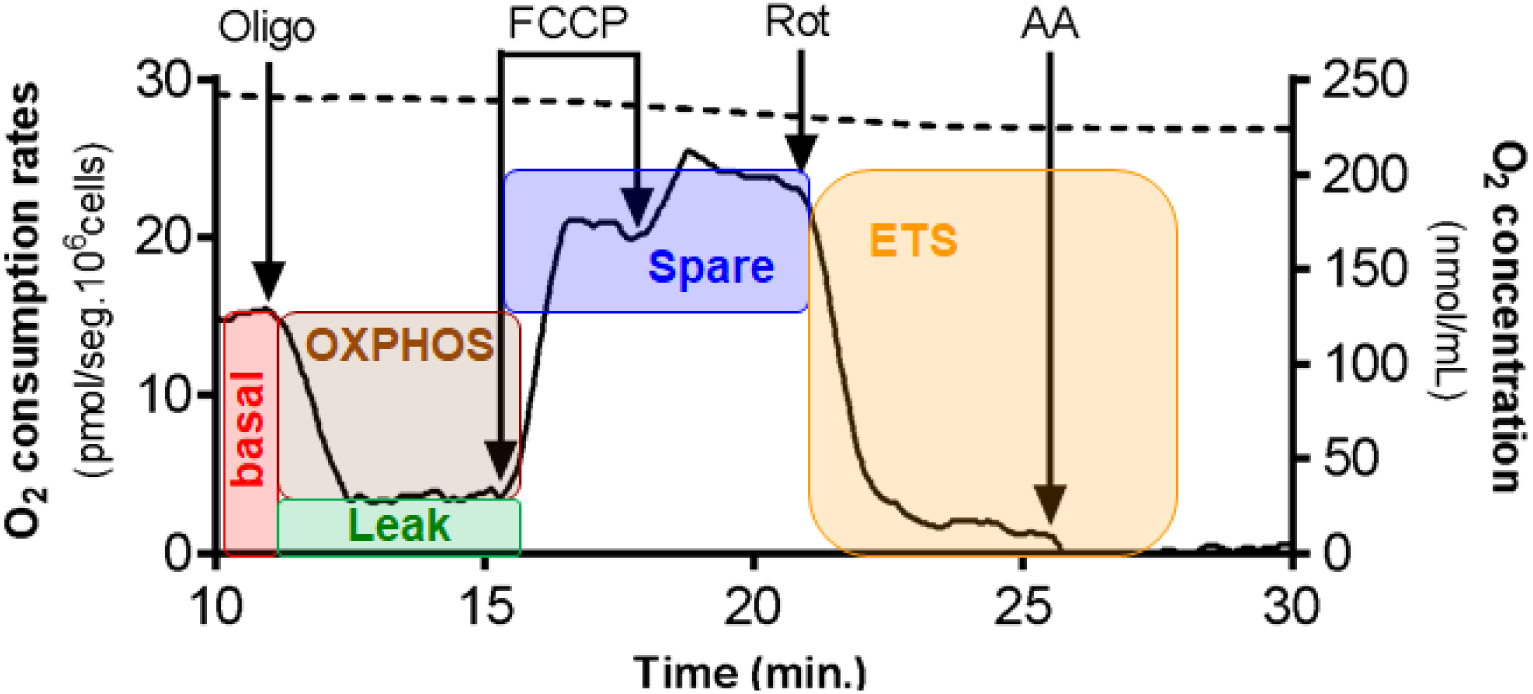
Representative oxygen consumption trace of intact S2 cells in Schneideŕs media. Respiratory rates were determined using 2 x 10^6^ S2 cells at 7^th^ day of culture at 26°C in Schneideŕs medium without FBS. Additions of OXPHOS modulators are represented with arrows, and their concentrations were the following: 250 ng/mL oligomycin (oligo), 300 nM FCCP (each injection), 100 nM rotenone (rot), 0,5 μg/mL antimycin a (AA). The solid line refers to oxygen consumption rates (left y axis), while the dashed line refers to oxygen tension in the chamber (right y axis). The five mitochondrial metabolic states (Basal, OXPHOS, Proton Leak, Spare Respiratory Capacity, Maximal ETS) were depicted as colored boxes within the oxygen consumption rate trace.

Basal, reflects the oxygen consumption rate of the cells in their culture condition, without any type of pharmacological interference; OXPHOS reflects the oxygen consumption rates linked to ATP synthesis and is mathematically defined by the difference between the basal respiratory rates from the oxygen consumption rates in the presence of oligomycin; Leak: reflects the oxygen consumption rates associated to proton leak through the inner mitochondrial membrane, since ATP synthase is blocked by oligomycin, and is defined by the difference between the respiratory rates under oligomycin treatment from those under antimycin A treatment; Spare respiratory capacity: reflects the increase of oxygen consumption rates from the basal state to the maximum respiratory rates produced by the addition of FCCP; Electron transport system (ETS): reflects the oxygen consumption rates produced after uncoupling by FCCP treatment and is mathematically defined by the difference between maximal respiratory rates produced by FCCP and the residual oxygen consumption after inhibition of complex III by the addition of antimycin A. In addition, to determine the potential effects of cell growth and temperature on the OXPHOS coupling, we calculated two flux control ratios (FCR) values of all the groups as described in the literature (Pesta and Gnaiger, 2012): the leak/ETS (L/E) and the OXPHOS/ETS (P/E) flux control ratios. The L/E ratios reflect an indirect assessment of OXPHOS coupling through altered proton leak relative to the ETS respiratory rates and span from 0 (fully coupled) to 1 (fully uncoupled) value. The P/E ratios represent the limiting effect of the mitochondrial phosphorylation system, which comprises the ATP synthase, the phosphate carrier, and the adenine nucleotide translocase, on respiratory rates relative to the ETS. P/E values span from minimal values of 0 (total inability to perform OXPHOS due to the maximal limitation of the phosphorylation system) to 1 (full OXPHOS capacity due to the maximal contribution of the phosphorylation system).

### 2.5. Data processing

The respirometry dataset was first parsed into a “tidy” long-format data frame to facilitate factorial analysis. All subsequent processing and statistical analyses were conducted using three AI tools: PerplexityPro (https://www.perplexity.ai/), Gemini Pro 2.5 (https://gemini.google.com/app?hl=pt-BR), and Formulabot (https://www.formulabot.com/), all using a standardized prompt (see Supplementary Data 1). To ensure robustness, outputs were systematically compared. When significant results were concordant, PerplexityPro data was presented. In cases of discrepancy, we adopted a conservative “majority rules” approach: a finding was only reported as statistically significant if at least two of the three platforms confirmed significance.

The complex multi-level headers were resolved into a structure containing three primary factors and one response column:

Time: A categorical factor with two levels (Early, Late). Temperature: A categorical factor with two levels (26°C, 28°C).

Group: A single categorical factor with four levels, created by combining the two main factors (Early-26°C, Early-28°C, Late-26°C, Late-28°C). This factor is used exclusively for *post-hoc* pairwise comparisons to analyze significant interactions.

Value: A continuous variable containing the measured data for each dependent variable.

Analyses were performed separately for the seven dependent variables, which were organized into two distinct functional groups: Group 1 (Respirometry: Basal, OXPHOS, Proton Leak, Spare Respiratory Capacity, Maximal ETS) and Group 2 (Flux Control Ratios: L/E, P/E).

### 2.6. Statistical analysis workflow

For each of the seven dependent variables, a rigorous three-step analytical workflow was executed. All statistical tests were performed with a significance level (alpha) of α = 0.05. Before primary analysis, the data for each variable were subjected to two assumption tests:

Normality: The Shapiro-Wilk test was used to assess the normality of the data within each of the four experimental groups. The null hypothesis (H_0_) is that the data are normally distributed. A p-value less than 0.05 indicates a violation of this assumption (Mishra et al., 2019).

Homogeneity of Variance (Homoscedasticity): Levene’s test was used to assess the equality of variances across all four experimental groups. The null hypothesis (H_0_) is that the variances are equal. *p*<0.05 indicates that the variances are not homogeneous (heteroscedasticity). The choice of the primary statistical test was conditional upon the results of the assumption testing:

Parametric path: If the data met both assumptions (all Shapiro-Wilk tests and the Levene’s test resulted in *p*>0.05), a Two-Way Analysis of Variance (ANOVA) was performed. The analysis was conducted using an Ordinary Least Squares (OLS) model with the formula Value ∼ C(Time) * C(Temperature), which assesses the main effect of Time, the main effect of Temperature, and the Time *vs.* Temperature interaction effect.

Non-parametric path: If the data violated either assumption (*p*<0.05 on any test), a robust non-parametric test was performed. The more powerful and modern Aligned Rank Transform (ART) ANOVA was selected. This method allows for the analysis of main effects and interactions in a non-parametric factorial design (Feys, 2016). If a significant main effect or interaction effect (p < 0.05) was detected in the primary analysis, a *post-hoc* test was performed to identify specific pairwise differences between the four experimental groups.

For Two-Way ANOVA: A Tukey’s Honestly Significant Difference (HSD) test was performed. As standard Python statsmodels packages do not directly compute Tukey’s HSD for interaction terms (statsmodels, n.d.), a valid workaround was employed: the 4-level Group factor (e.g., “Early-26°C") was used as the input for the statsmodels.stats.multicomp.pairwise_tukeyhsd function. This provides a simultaneous, corrected comparison of all four experimental conditions (statsmodels, n.d.). We also employed Šídák’s multiple comparisons test for Group 2 (Flux Control Ratios) comparisons by using the GraphPad Prism software version 8.0 for Windows (GraphPad Software, USA).

For ART ANOVA: Pairwise comparisons were conducted on the aligned ranks using a Dunn’s test with p-values adjusted for multiple comparisons using the Bonferroni correction. This was applied to the 4-level Group factor to dissect the significant effects. All graphs were generated by GraphPad Prism software version 8.0 for Windows (GraphPad Software, USA), displaying the ± standard error of the mean (SEM) of values. Significance notations derived from the *post-hoc* tests are displayed as compact letters above each bar (*p*<0.05).

## 3. Results

### 3.1. Experimental outline

We used the Schneider S2 *Drosophila* embryonic cell line (S2) to examine how variations in cell growth phase and culture temperature influence mitochondrial bioenergetic capacity and efficiency. High-resolution respirometry was used to measure five mitochondrial respiratory states: basal respiration, oxidative phosphorylation (OXPHOS), proton leak, spare respiratory capacity, and maximal electron transport system (ETS) capacity. Measurements were carried out at two culture temperatures (26°C and 28°C) and at two time points corresponding to early (E) and late (L) growth phases. To evaluate how temperature and growth phase affected mitochondrial physiology, we applied a series of data processing and statistical analyses as described in the Methods section. These analyses tested both main effects and the interaction between temperature and growth phase as outlined in Figure 2. Data were analyzed across seven dependent variables, grouped into two functional categories:

**Figure 2:**
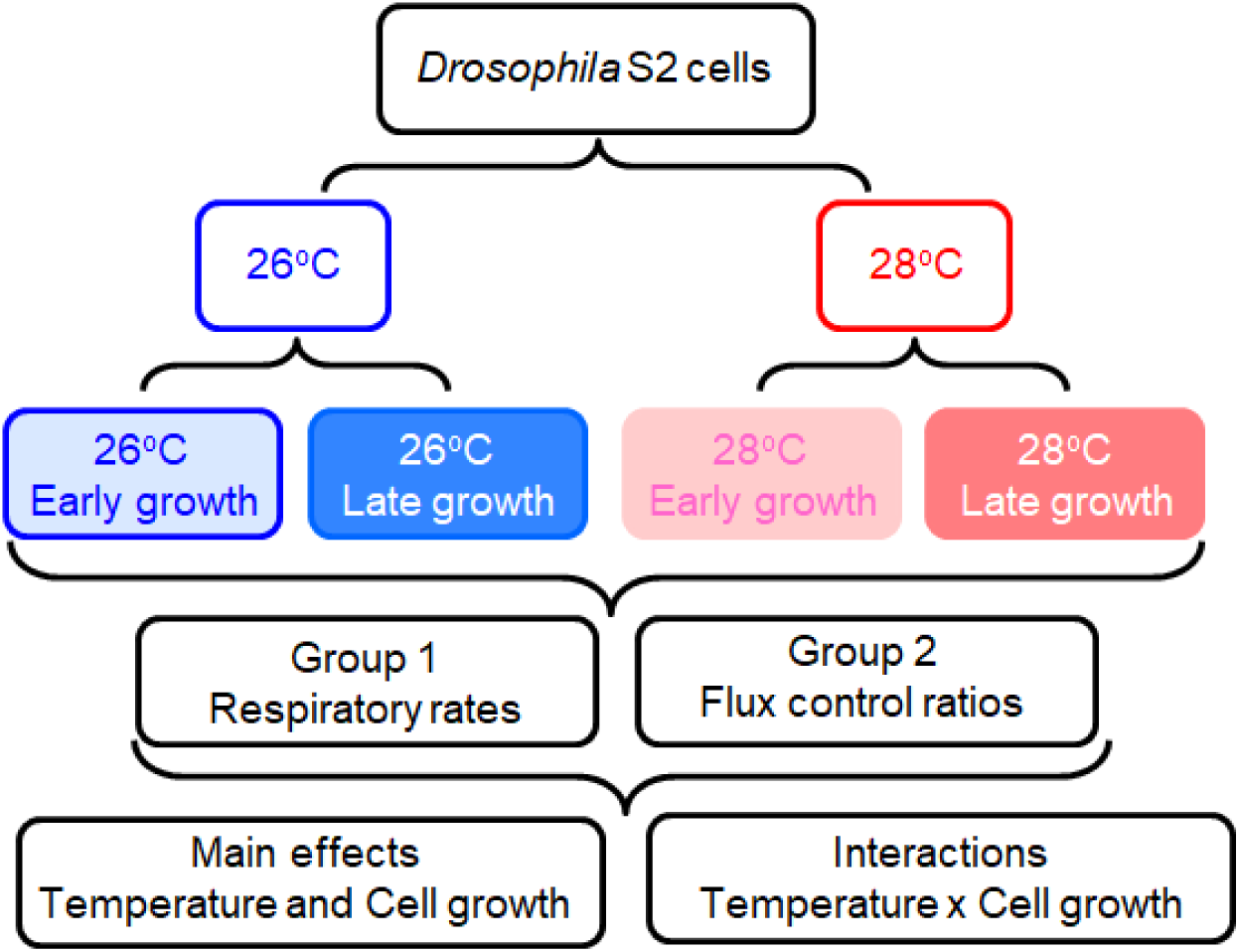
Experimental outline for mitochondrial metabolic analysis in *Drosophila* S2 cells. *Drosophila* S2 cells were cultured at two distinct temperatures (26°C, blue, and 28°C, red) and assessed at two stages of cell growth (early growth at the 2^nd^ day of culture, light colors, and late growth at the 7^th^ day, dark colors). Cellular respirometry analysis was then performed on all four resulting experimental conditions to determine five mitochondrial metabolic states (group 1) as well as two flux control ratios (group 2). Subsequent analysis focuses on the main effects of temperature and cell growth, as well as their interactive effects on these parameters.

Group 1: Respirometry variables (basal, OXPHOS, proton leak, spare respiratory capacity, maximal respiration)

Group 2: Flux Control Ratios (L/E, P/E).

### 3.2. Assessment of S2 cell growth and respiratory rates

In order to provide context for our metabolic analyses, we first characterized the proliferation of the Schneider S2 cell line at 26°C and 28°C over nine days (Figure 2). Cells were seeded at 8.8 × 10³ cells/well on day 0, and their growth was monitored throughout the period. Consistent with Schneider’s original 1972 study, we observed a lag phase of minimal growth during the first two days, followed by robust proliferation from day four onward, reaching a plateau near days eight to nine (Schneider, 1972). A study reviewing strategies for heterologous gene expression in *Drosophila* S2 cells also examined the influence of bioprocess parameters on cell growth. The authors reported that S2 cells achieved high maximum specific growth rates and maximum cell concentrations at temperatures between 24°C and 28°C. Notably, no significant differences in these growth parameters were observed within this specific temperature range (Moraes et al., 2012).

Our primary objective was to investigate the impact of a minor (2°C) temperature increase—a shift shown to affect several cellular processes in various systems (Jalal et al., 2015; Rao and Engelberg, 1965; Shao-Hua et al., 1998). The routine culture temperature for S2 cells varies depending on the cell provider, ranging from approximately 23°C to around 27°C, although studies have also assessed this cell line at slightly higher temperatures (Bouchet et al., 2014; “Drosophila S2 Cells in Schneider’s Medium 1 mL | Contact Us | Gibco^TM^ | thermofisher.com,” n.d.; “Schneider’s Drosophila Line 2 [D. Mel. (2), SL2] - CRL-1963 | ATCC,” n.d.). Growth curves at 26°C and 28°C showed no statistically significant differences in maximum specific growth rates (0.59 *vs.* 0.51 x day^-1^, respectively) or maximum cell concentrations (3.58 *vs*. 3.53 x 10^6^ cells/well, respectively) (Figure 3). This consistent growth behavior enabled the selection of comparable time points for respirometry measurements at both temperatures: day 2 representing the early growth phase and day 7 for the late exponential growth phase. Slight temperature variations such as these have been reported to modulate energy metabolism by altering mitochondrial efficiency and metabolic rate, without necessarily impacting cell proliferation markedly within this narrow range (Klepsatel et al., 2019; Moraes et al., 2012). Overall, this approach provides a controlled framework to evaluate temperature-dependent bioenergetic changes independent of major growth rate alterations.

**Figure 3:**
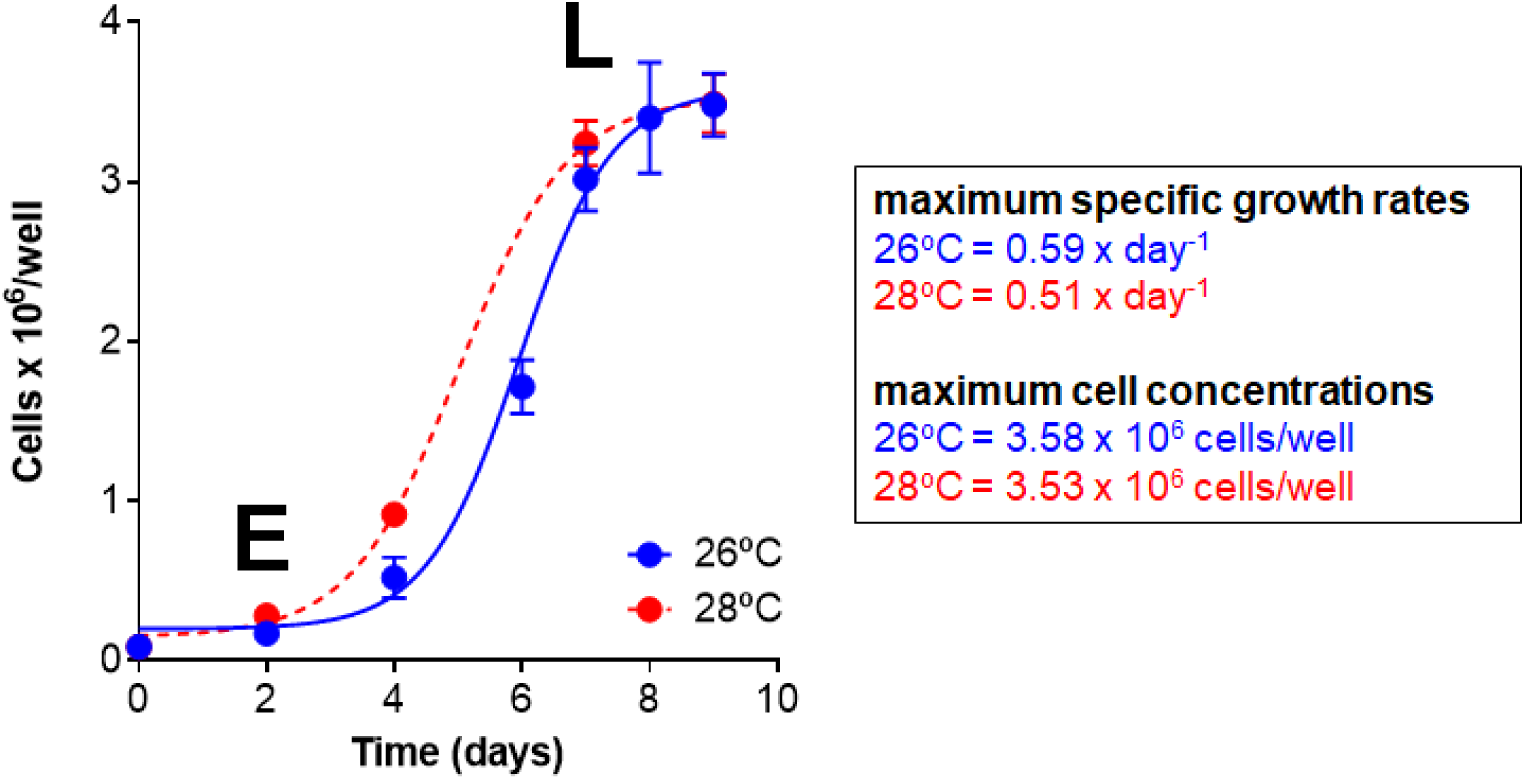
S2 cell growth in Schneider’s media at two different temperatures. An aliquot corresponding to 8.8 x 10^3^ cells was seeded/well and maintained in Schneider medium supplemented with 10% SFB for up to 9 days at 26°C (blue circles) and 28°C (red circles). Only adherent cells were used throughout this study. Cell counts were performed through the Neubauer chamber using Trypan blue as a cell viability stain. Data are expressed as mean ± S.E.M. of three different experiments carried out at least in triplicate. The early (E, at 2 days of culture) and late (L, at 7 days of culture) times investigated throughout this study are defined in the graph. Sigmoidal dose-response non-linear regression analyses were performed for both groups and are presented at 26°C (solid blue line) and 28°C (dashed red line).

### 3.3. Basal, OXPHOS and leak respiration

Mitochondrial physiology in intact S2 cells was evaluated using high-resolution respirometry. Figure 1 provides a representative trace from an experiment (7-day culture at 26°C), illustrating the protocol used to define the mitochondrial metabolic states. The protocol involved the sequential addition of mitochondrial modulators (oligomycin, FCCP, rotenone and antimycin A) to isolate specific respiratory states, as detailed in the methods section. We then compared the five respiratory states across the four conditions (2 temperatures x 2 growth phases) and the following sections show the results for each mitochondrial bioenergetic parameter.

Basal respiration assesses the oxygen consumption of intact cells using available nutrients in the culture medium. This mitochondrial bioenergetic parameter reflects aerobic energy demand under standard physiological conditions and depends on factors such as nutrient supply, cellular viability, proliferative status, and mitochondrial efficiency (Doerrier et al., 2018; Gnaiger, 2020; Pesta and Gnaiger, 2012). A Two-Way ANOVA with Tukey’s HSD *post-hoc* test was applied to compare the four experimental groups. Significant main effects of time, temperature, and interaction were observed (Table S1). *Post-hoc* analysis revealed that basal respiration in the Late-28°C group was significantly higher than in all other groups (*p*<0.001) (Figure 4A, Table S2). No other differences were detected. Thus, the strong interaction between time and temperature was primarily driven by the Late-28°C condition, where the combined effects exceeded the additive outcome of each factor alone, indicating a synergistic elevation in basal respiratory rate.

**Figure 4:**
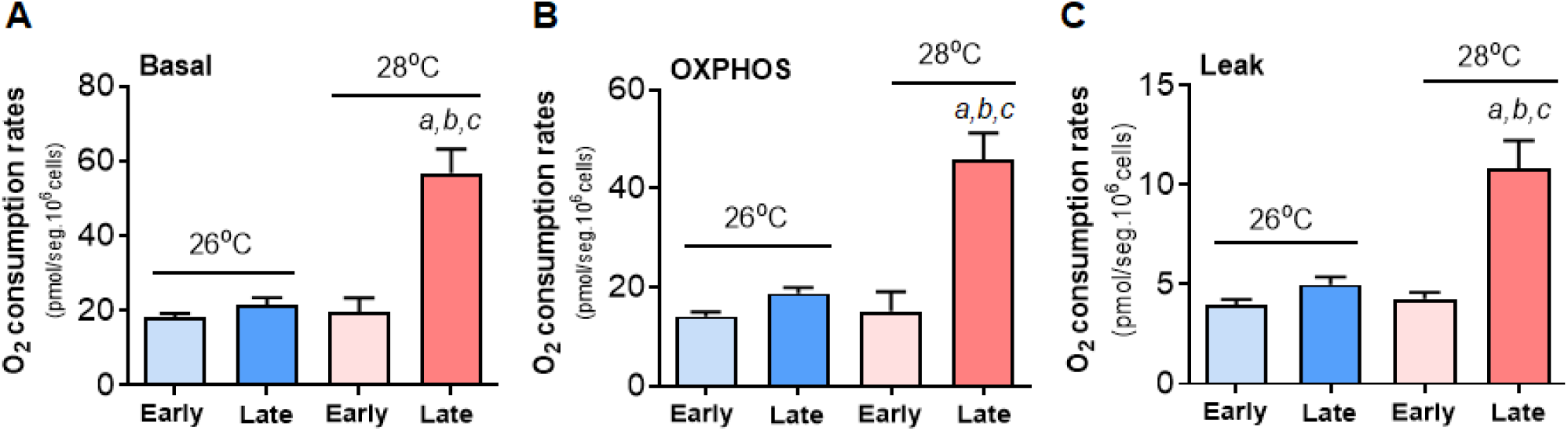
Effect of time and temperature on basal, OXPHOS and leak respiration. Oxygen consumption rates (OCR, O₂ flow per cell; pmol/sec·10⁶ cells) in the basal (A), OXPHOS (B) and leak (C) states were determined in S2 cells at early (2^nd^ day, light bars, n=6) and late (7^th^ day, dark bars, n=3-22) time growth at 26°C (blue bars, n=6-22) and at 28°C (red bars, n=3-6). All data are presented as the mean ± SEM from at least three independent experiments. Statistical comparisons between the four experimental groups were performed using a Two-Way ANOVA, followed by Tukey’s HSD *post-hoc* test for multiple comparisons. In all figures statistical differences were identified as following: *a* Early-26°C *vs.* Late-28°C, *p*<0.001; *b* Early-28°C *vs.* Late-28°C, *p*<0.001; *c* Late-26°C *vs*. Late-28°C, *p*<0.001).

We next analyzed the ATP-linked respiratory rate, which represents the fraction of basal respiration coupled to ATP synthesis through oxidative phosphorylation (OXPHOS) (Doerrier et al., 2018; Gnaiger, 2020; Pesta and Gnaiger, 2012). In this state, mitochondria consume oxygen at a high rate driven essentially by the energy demand, primarily the availability of ADP and inorganic phosphate, which stimulate ATP synthase to phosphorylate ADP into ATP using the proton gradient generated by the electron transport system. Following oligomycin addition, the reduction in oxygen consumption reflected the OXPHOS contribution, which represented 72–83% of total respiration across all conditions—indicative of well-coupled mitochondrial oxygen consumption in S2 cells.

A Two-Way ANOVA confirmed significant main effects for time, temperature, and a significant time *vs.* temperature interaction on OXPHOS respiration (Table S3). This interaction proved to be synergistic: *post-hoc* tests showed that, similar to basal respiration, only the Late-28°C group exhibited a significant increase in OXPHOS capacity compared to all other groups (all *p*<0.001; Figure 4B, Table S4). This suggests the substantial rise in basal respiration in the Late-28°C group is primarily driven by this enhanced OXPHOS capacity, reflecting a greater allocation to mitochondrial ATP production.

Leak respiration measures oxygen consumption required to offset proton leakage across the inner mitochondrial membrane and was assessed after oligomycin addition, with non-mitochondrial oxygen consumption subtracted following antimycin A treatment (Pesta and Gnaiger, 2012). In this state, the electron transport system remains active, but its sole function is to pump protons to maintain the mitochondrial membrane potential (ΔΨ_m_) against the leak. In living cells, this leak state is dynamically regulated by two main factors: the mitochondrial membrane potential (the driving force) and the proton conductance (the “leakiness") of the inner mitochondrial membrane (IMM). Therefore, an observed change in leak respiration indicates a change in either this driving force or the IMM physical properties. A Two-Way ANOVA with Tukey’s *post-hoc* test (Table S5) indicated significant main effects for time, temperature, and their interaction. Consistent with previous analyses, only the Late-28°C group showed a marked increase in leak respiration compared with all other groups (*p*<0.001; Figure 4C, Table S6). However, leak respiration contributed only about 22% of total basal oxygen consumption, underscoring that the heightened basal metabolism at Late-28°C was predominantly due to enhanced OXPHOS activity rather than increased proton leak.

### 3.4. Spare respiratory capacity and maximal respiration

Spare respiratory capacity is a critical measure of mitochondrial physiology, defined as the difference between maximal respiration and basal respiration. Experimentally, the maximal respiration rate is determined by adding a proton ionophore (such as FCCP), which uncouples oxygen consumption from ATP synthesis. By collapsing the ΔΨ_m_, the ionophore forces the electron transport system to operate at its maximum flux. The spare respiratory capacity is, therefore, the value calculated from this measurement: the difference between the maximal, ionophore-induced oxygen consumption rate and the cell’s basal respiration rate (Doerrier et al., 2018; Gnaiger, 2020; Pesta and Gnaiger, 2012). This “spare” capacity represents the bioenergetic reserve the cell can engage to produce ATP when faced with sudden increases in energy demand or acute stress. A high spare respiratory capacity, therefore, indicates metabolic robustness, allowing cells to survive insults, whereas a low spare respiratory capacity indicates the cell is operating near its bioenergetic limit and indicates vulnerability to dysfunction and death (Choi et al., 2009; Nicholls, 2009; Yadava and Nicholls, 2007).

A Two-Way ANOVA revealed strongly significant main effects of time, temperature, and their interaction (Table S7). The *post-hoc* analysis highlighted the powerful effect of these factors on spare respiratory capacity (Figure 5A, Table S8). While the Late-28°C group exhibited the most dramatic increase in spare respiratory capacity (*p*<0.001 vs. all other groups), a finding unique to this assessment—and not observed in basal, OXPHOS or leak respiration—was the significant increase in the Late-26°C group compared to the Early-26°C group. This distinction is critical as it demonstrates that as cells mature, they significantly enhance their metabolic reserve to handle acute stress, even without the added stimulus of temperature. This unique, spare respiratory capacity-specific finding suggests that the enhancement of maximal respiration outpaces any changes in the other metabolic states. This points toward an increase in functional respiratory components, likely via mitochondrial biogenesis, as the primary underlying mechanism, rather than just an increased supply of substrates.

**Figure 5:**
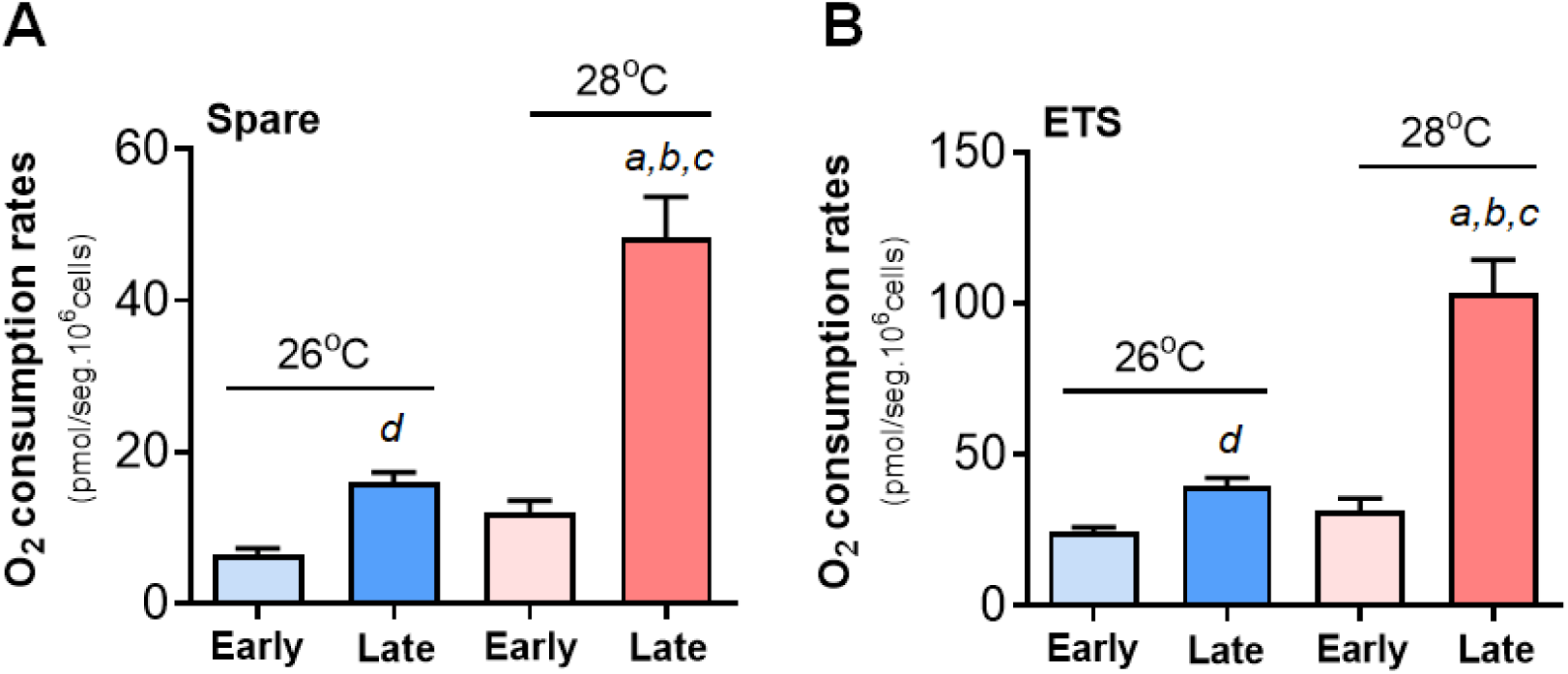
Effect of time and temperature on spare respiratory capacity and maximal respiration. Oxygen consumption rates (OCR, O₂ flow per cell; pmol/sec·10⁶ cells) in the spare respiratory capacity (A) and in the maximal respiratory rates (ETS) (B) states were determined in S2 cells at early (2^nd^ day, light bars, n=6) and late (7^th^ day, dark bars, n=3-18) time growth at 26°C (blue bars, n=6-18) and at 28°C (red bars, n=3-6). All data are presented as the mean ± SEM from at least three independent experiments. Statistical comparisons between the four experimental groups were performed using a Two-Way ANOVA, followed by Tukey’s HSD *post-hoc* test for multiple comparisons. In all figures statistical differences were identified as following: *a* Early-26°C *vs.* Late-28°C, *p*<0.0001; *b* Early-28°C *vs.* Late-28°C, *p*<0.0001; *c* Late-26°C *vs*. Late-28°C, *p*<0.0001; *d* Early-26°C *vs.* Late-26°C, *p*=0.0042 for figure A and *p*<0.03 for figure B).

The final mitochondrial metabolic state assessed was electron transfer system (ETS) capacity. This refers to the maximal rate of oxygen consumption when the ETS is operating at its theoretical maximum, entirely independent of ATP synthesis. The ETS capacity is experimentally achieved by adding proton ionophores (such as FCCP or DNP). These compounds dissipate the ΔΨ_m_ which in turn removes the proton back-pressure on the ETS, allowing electrons to flow unrestrictedly to oxygen (Doerrier et al., 2018; Gnaiger, 2020; Pesta and Gnaiger, 2012).

The effects of culture time and temperature on maximal respiratory rates largely mirrored those observed for spare respiratory capacity, demonstrating significant main effects of time, temperature, and a robust interaction between these factors (Table S9). *Post-hoc* analysis confirmed the combined impact, revealing the most marked increase in the Late-28°C group (Figure 5A, 5B; Table S10). Notably, cell maturation alone (comparing Late-26°C to Early-26°C) significantly enhanced both maximal ETS respiration (*p*=0.026) and spare respiratory capacity. This suggests that maturation enhances mitochondrial respiratory capacity, independent of thermal stress. This specific pattern, distinct from other respiratory states, points to an increase in mitochondrial respiratory capacity—likely via biogenesis or augmented TCA cycle/ETS complex activities—rather than simply greater substrate availability.

Together, these data demonstrate a coordinated enhancement of mitochondrial physiology driven by both maturation and thermal adaptation. The result is an elevated metabolic reserve capacity, boosted maximal respiration, and strengthened cellular bioenergetic flexibility to meet acute energy demands.

### 3.5. Flux control ratios

In the last set of analyses, we determined the effects of time of culture and temperature on the flux control ratios (FCR) of S2 cells as described in the literature (Doerrier et al., 2018; Gnaiger, 2020; Pesta and Gnaiger, 2012). FCRs, specifically the leak/ETS (L/E) and OXPHOS/ETS (P/E) coupling control ratios, are essential quantitative measures in mitochondrial physiology used to evaluate respiratory states normalized to the ETS capacity. The L/E ratio represents the fraction of respiration attributable to proton leak relative to maximal ETS capacity, ranging from 0 for a fully coupled system (minimal leak) to 1 for a fully uncoupled system (maximal leak), thus serving as an indirect index of OXPHOS coupling efficiency. The P/E ratio reflects the limiting influence of the mitochondrial phosphorylation system—which includes ATP synthase, phosphate carrier, and adenine nucleotide translocase—on the OXPHOS respiratory rate relative to ETS capacity; values span from 0, indicating complete inhibition or incapacity of oxidative phosphorylation due to phosphorylation system limitations, to 1, indicating maximal OXPHOS capacity with full contribution from the phosphorylation system. Together, these ratios provide biologically and metabolically significant insights into mitochondrial coupling efficiency, energy conversion capacity, and functional constraints, enabling standardized comparison across samples independent of mitochondrial content or experimental variability.

A Two-Way ANOVA revealed a significant main effect of time on the L/E ratio, but no significant effect of temperature or interaction between time and temperature (Table S11). *Post-hoc* analysis showed a significant difference only between Early-26°C and Late-28°C groups, with no other differences observed for L/E (Figure 6A, Table S12). For the P/E ratio, the Two-Way ANOVA indicated significant main effects of time and a significant time by temperature interaction (Table S13). Consistent with L/E results, *post-hoc* tests revealed a significant difference only between Early-26°C and Late-26°C groups (Figure 6B, Table S14).

**Figure 6:**
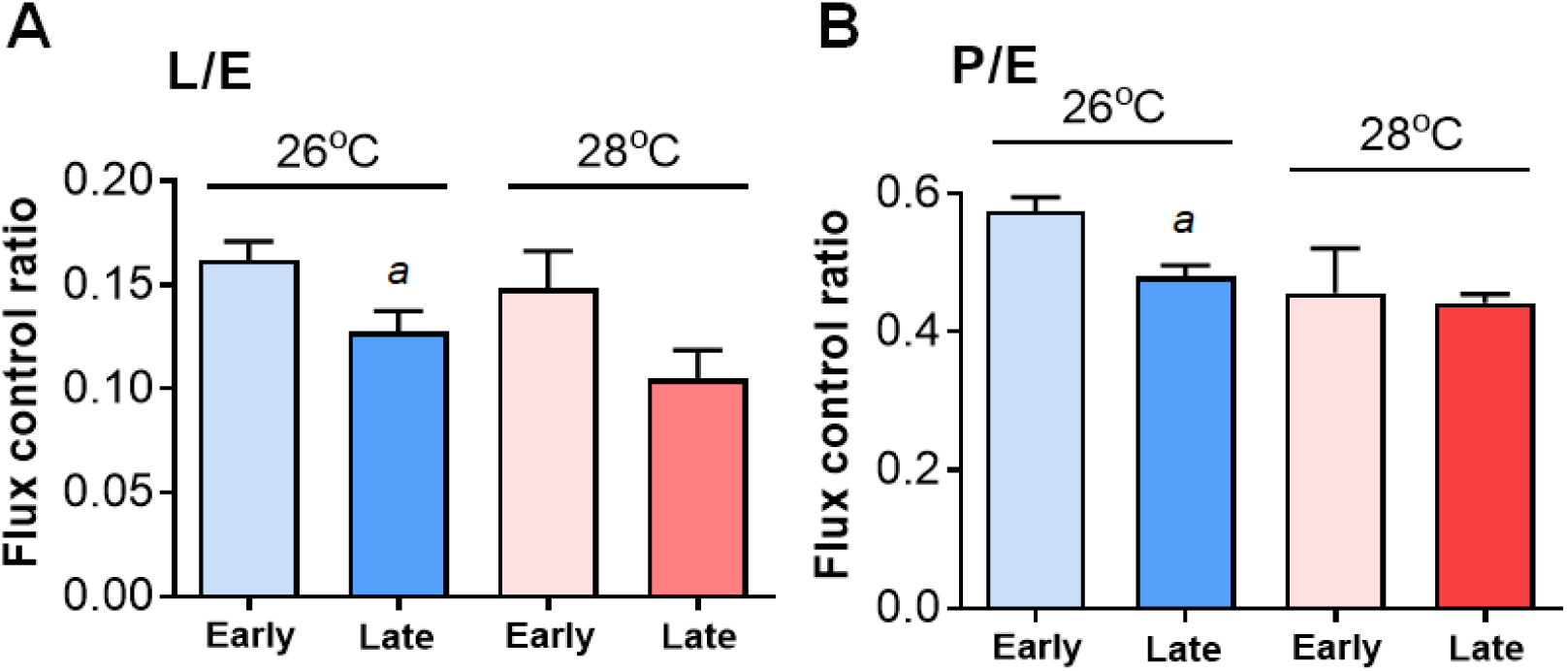
Effect of time and temperature on flux control ratios. (A) Leak/ETS (L/E) and (B) OXPHOS/ETS (P/E) flux control ratios were determined in S2 cells at early (2^nd^ day, light bars, n=6) and late (7^th^ day, dark bars, n=3-18) time growth at 26°C (blue bars, n=6-18) and at 28°C (red bars, n=3-6). All data are presented as the mean ± SEM from at least three independent experiments. Statistical comparisons between the four experimental groups were performed using a Two-Way ANOVA, followed by Tukey’s HSD *post-hoc* test (A) or Šídák’s multiple comparisons test (B) for multiple comparisons. In all figures statistical differences were identified as following: (A and B) *a* Early-26°C *vs.* Late-26°C, *p*<0.05.

These findings suggest that the effects of culture time on both FCRs (L/E and P/E) are more pronounced at lower temperature (26°C). This may be due to a disproportionate increase of ETS capacity relative to both leak and OXPHOS states at this temperature. Biologically, this means that the coupling efficiency between respiration and ATP synthesis is generally maintained but significantly reduced only at 26°C during late culture times compared to early times.

## 4. Discussion

The findings presented here demonstrate that both progression of cell growth and slight temperature increases significantly activate mitochondrial physiology, mostly through increased mitochondrial maximal and spare respiratory capacities. The observed changes in mitochondrial physiology presented here strongly align with reported heat stress-induced mitochondrial biogenesis and OXPHOS remodeling. In addition, given that cell proliferation remained largely consistent across temperature groups, the increased energy demand appears to be driven by cell enlargement rather than division. Enhanced bioenergetic capacity with preserved efficiency would allow larger cells to sustain energy homeostasis while suggesting the existence of an optimal cell size for maximal mitochondrial efficiency (Miettinen et al., 2017; Miettinen and Björklund, 2017). This regulation is not only a response to increased energy demand but also reflects a fundamental adaptation of mitochondrial physiology to cellular and environmental cues. The observed increases in mitochondrial bioenergetic capacity are consistent with previous studies showing that mitochondrial metabolism is highly sensitive to both developmental and thermal stimuli (Gerber et al., 2021; Liu and Brooks, 2011; Lu et al., 2023a; Roussel and Voituron, 2020; Sokolova, 2023; Thoral et al., 2025).

Mitochondria are central to cellular energy homeostasis, and their physiology is profoundly influenced by temperature. Incubation at higher temperatures has been shown to increase mitochondrial respiration rates and ATP synthesis, albeit at the cost of reduced coupling efficiency (Roussel and Voituron, 2020; Thoral et al., 2025). This phenomenon is particularly relevant in ectothermic organisms, in which temperature directly influences mitochondrial membrane fluidity and enzyme activity, thereby modulating overall metabolic output. In our study, slight temperature increases likely enhanced mitochondrial oxygen consumption, primarily by elevating spare and maximal respiratory capacities. This finding supports the idea that mitochondria can rapidly adjust to thermal challenges by upregulating their bioenergetic machinery, consistent with evidence reported in the literature (Hafen et al., 2018; Liu and Brooks, 2011; Maunder et al., 2024; Roussel and Voituron, 2020; Thoral et al., 2025).

The mitochondrial adaptations observed in the present study, particularly the disproportionate increase in mitochondrial maximal and spare respiratory capacities over culture time and varying temperatures, align well with reported literature on heat stress-induced mitochondrial biogenesis and remodeling. Several studies show that mild heat stress induces mitochondrial biogenesis through activation of key regulators such as AMPK, SIRT1, and PGC-1α. For instance, mild heat stress in C2C12 myotubes increased PGC-1α protein levels, oxidative phosphorylation proteins, and mitochondrial DNA copy number, mediated by the AMPK-SIRT1-PGC-1α signaling pathway (Liu and Brooks, 2011). This pathway plausibly explains your observed increase in mitochondrial components with culture time and temperature, suggesting that mild heat promotes mitochondrial biogenesis rather than just physiological adaptation (Liu and Brooks, 2011). Repeated heat exposure in human skeletal muscle also leads to increased expression of mitochondrial electron transport proteins and enhanced respiratory capacity, further supporting the role of mild heat as a positive modulator of mitochondrial adaptation (Hafen et al., 2018). On the other side, acute or severe heat stress can have detrimental effects, disrupting mitochondrial physiology, reducing membrane potential, and inducing apoptosis (Lu et al., 2023b; Von Schulze and Geiger, 2022). Our data, showing preserved coupling efficiency at 28°C despite large increases in mitochondrial parameters, is consistent with beneficial mild heat adaptation, while reduced coupling at 26°C may reflect early imbalances between ETS capacity and coupling components. In this sense, it seems that mitochondrial components involved in electron transfer within the ETS complexes increase more rapidly than components involved in proton leak (such as mitochondrial carriers), ATP synthase subunits, adenine nucleotide translocator (ANT) and the phosphate carrier. Despite increased mitochondrial biogenesis or reduced mitophagy suggestive of higher mitochondrial content—especially evident at 28°C with large increases in all measured bioenergetic parameters—the coupling control remains preserved at the higher temperature. The reduced coupling observed at 26°C likely results from the disproportionate rise in ETS capacity rather than reductions in leak or OXPHOS states. Additionally, the increase in substrate supply supporting electron transport, driven by culture time and temperature, is more pronounced in the Late-28°C group than in the Late-26°C group. This substrate supply enhancement boosts bioenergetic capacity without compromising efficiency and is shaped by the positive interaction between slight temperature increases and cell growth.

Cell growth, rather than proliferation, emerges as a key determinant of mitochondrial bioenergetic capacity (Chacko et al., 2025). Mitochondrial DNA copy number and network volume scale proportionally with cell volume, ensuring that larger cells maintain adequate mitochondrial mass and metabolic output (Seel et al., 2023). However, mitochondrial metabolic activity exhibits a nonlinear relationship with cell size: functionality is highest in intermediate-sized cells, while very large or small cells show reduced mitochondrial membrane potential and metabolic efficiency (Miettinen et al., 2017). This nonlinear scaling ensures that mitochondrial bioenergetic capacity matches the metabolic needs of the cell, regardless of whether growth is driven by proliferation or hypertrophy (Chacko et al., 2025; Seel et al., 2023).

The increase in cell size, rather than cell proliferation, may represent a primary energy demand process that drives the upregulation of mitochondrial bioenergetic capacity. Larger cells require more ATP to maintain cellular homeostasis, and this demand is met by expanding mitochondrial networks and enhancing OXPHOS. The nonlinear scaling of mitochondrial metabolic activity with cell size has been documented in both proliferating and non-proliferating cells, highlighting the importance of cell size as a regulator of mitochondrial physiology (Chacko et al., 2025; Miettinen et al., 2017; Miettinen and Björklund, 2017; Seel et al., 2023). This scaling ensures that mitochondrial bioenergetic capacity is matched to the metabolic needs of the cell, regardless of whether growth is driven by proliferation or hypertrophy. Nevertheless, cell proliferation also imposes substantial energetic demands, requiring not only increased bioenergetic capacity but also greater efficiency, a process that appears to involve mitochondrial fusion as a mechanism to support these demands (Yao et al., 2019).

## 5. Conclusions

In summary, our results indicate that slight temperature increases and cell growth promote mitochondrial adaptations that enhance bioenergetic capacity without impairing coupling efficiency, as long as thermal stress remains within physiological limits. The scaling of mitochondrial bioenergetic capacity in S2 cells under these conditions likely reflects the activation of a mitochondrial biogenesis program that maintains efficiency while expanding mitochondrial bioenergetic capacity. These findings have broad implications for understanding mitochondrial plasticity in response to environmental and developmental cues. They may inform strategies to optimize mitochondrial physiology in several biological contexts, such as aging, metabolic disorders, and adaptation to different stress signals.

## CRediT authorship contribution statement

Rodiésley S. Rosa: Investigation, Methodology, Data Curation, Writing – Review & Editing. Ana Paula M. Mendonça: Conceptualization, Supervision, Formal Analysis, Writing – Original Draft, Review. Matheus P. Oliveira: Conceptualization, Supervision, Formal Analysis, Writing – Original Draft, Review. Danielle B. Carvalho: Investigation, Data Collection, Data Analysis. Leonardo Vazquez: Supervision, Critical Review, Final Approval. Marcus F. Oliveira: Conceptualization, Funding Acquisition, Project Administration, Supervision, Writing – Review & Editing, Final Approval.

## Declaration of competing interest

The authors declare that they have no known competing financial interests or personal relationships that could have appeared to influence the work reported in this paper.

## Supporting information

AI prompt for statistical analyses

Full respirometry dataset

## Acknowledgements

We are grateful for the valuable comments and criticisms raised by the editor and the reviewers on the original version of the manuscript. This study was financed in part by the Coordenação de Aperfeiçoamento de Pessoal de Nível Superior – Brasil (CAPES) – Finance Code 001, by the Conselho Nacional de Desenvolvimento Científico e Tecnológico (CNPq) [grant number: 308629/2021-3] and Fundação de Amparo à Pesquisa do Estado de São Paulo [grant number: 2021/06711-2]. The funders had no role in study design, data collection and analysis, decision to publish, or preparation of the manuscript.

## Data availability

Original data will be made available upon request.

## Declaration of generative AI and AI-assisted technologies in the writing process

During the preparation of this work the author(s) used the AI language models Gemini Pro 2.5 and Perplexity Pro for two purposes: i) to carry out data processing and statistical analyses; ii) to enhance the English language. Description of data processing and statistical analyses are provided in the supplementary files. When revisions were needed, the models were instructed with the prompt: “Revise the following section in order to make it clearer and with an English-native speaker writing”. The authors retained full responsibility for the manuscript’s content, and all AI-assisted revisions were critically evaluated to confirm that the original scientific information was preserved with precision and correctness.

### Abbreviations

ADP: adenosine diphosphate;
ATP: adenosine triphosphate;
AA: antimycin A;
BSA: bovine serum albumin;
ETS: electron transport system;
FADH₂: flavin adenine dinucleotide (reduced);
FBS: fetal bovine serum;
FCCP: carbonyl cyanide p-(trifluoromethoxy)phenylhydrazone;
FCR: flux control ratio;
NADH: nicotinamide adenine dinucleotide (reduced);
OCR: oxygen consumption rate;
OXPHOS: oxidative phosphorylation;
S2: Drosophila melanogaster Schneider-2 cell line;
SEM: standard error of the mean.

## Notes

### Competing Interest Statement

The authors have declared no competing interest.

## References

1. Abrams, J.M., Lux, A., Steller, H., Krieger, M., 1992. Macrophages in Drosophila embryos and L2 cells exhibit scavenger receptor-mediated endocytosis. Proc Natl Acad Sci U S A 89, 10375–10379. 10.1073/pnas.89.21.10375

2. Bouchet, N., Jaillet, J., Gabant, G., Brillet, B., Briseño-Roa, L., Cadene, M., Augé-Gouillou, C., 2014. cAMP protein kinase phosphorylates the Mos1 transposase and regulates its activity: evidences from mass spectrometry and biochemical analyses. Nucleic Acids Res 42, 1117–1128. 10.1093/nar/gkt874

3. Chacko, L.A., Nakaoka, H., Morris, R.G., Marshall, W.F., Ananthanarayanan, V., 2025. Mitochondrial function regulates cell growth kinetics to maintain mitochondrial homeostasis. Curr Biol 35, 5278–5288.e4. 10.1016/j.cub.2025.09.046

4. Cherbas, L., Willingham, A., Zhang, D., Yang, L., Zou, Y., Eads, B.D., Carlson, J.W., Landolin, J.M., Kapranov, P., Dumais, J., Samsonova, A., Choi, J.-H., Roberts, J., Davis, C.A., Tang, H., van Baren, M.J., Ghosh, S., Dobin, A., Bell, K., Lin, W., Langton, L., Duff, M.O., Tenney, A.E., Zaleski, C., Brent, M.R., Hoskins, R.A., Kaufman, T.C., Andrews, J., Graveley, B.R., Perrimon, N., Celniker, S.E., Gingeras, T.R., Cherbas, P., 2011. The transcriptional diversity of 25 Drosophila cell lines. Genome Res 21, 301–314. 10.1101/gr.112961.110

5. Choi, S.W., Gerencser, A.A., Nicholls, D.G., 2009. Bioenergetic analysis of isolated cerebrocortical nerve terminals on a microgram scale: spare respiratory capacity and stochastic mitochondrial failure. Journal of Neurochemistry 109, 1179–1191. 10.1111/j.1471-4159.2009.06055.x

6. Chrétien, D., Bénit, P., Ha, H.-H., Keipert, S., El-Khoury, R., Chang, Y.-T., Jastroch, M., Jacobs, H.T., Rustin, P., Rak, M., 2018. Mitochondria are physiologically maintained at close to 50 °C. PLOS Biology 16, e2003992. 10.1371/journal.pbio.2003992

7. Doerrier, C., Garcia-Souza, L.F., Krumschnabel, G., Wohlfarter, Y., Mészáros, A.T., Gnaiger, E., 2018. High-Resolution FluoRespirometry and OXPHOS Protocols for Human Cells, Permeabilized Fibers from Small Biopsies of Muscle, and Isolated Mitochondria. Methods Mol Biol 1782, 31–70. 10.1007/978-1-4939-7831-1_3

8. Drosophila S2 Cells in Schneider’s Medium 1 mL | Contact Us | Gibco^TM^ | thermofisher.com [WWW Document], n.d. URL https://www.thermofisher.com/order/catalog/product/br/en/R69007 (accessed 11.6.25).

9. Elwell, C., Engel, J.N., 2005. Drosophila melanogaster S2 cells: a model system to study Chlamydia interaction with host cells. Cell Microbiol 7, 725–739. 10.1111/j.1462-5822.2005.00508.x

10. Feys, J., 2016. New Nonparametric Rank Tests for Interactions in Factorial Designs with Repeated Measures. Journal of Modern Applied Statistical Methods 15, 78–99. 10.22237/jmasm/1462075500

11. Gerber, L., Clow, K.A., Gamperl, A.K., 2021. Acclimation to warm temperatures has important implications for mitochondrial function in Atlantic salmon (Salmo salar). J Exp Biol 224, jeb236257. 10.1242/jeb.236257

12. Gnaiger, E., 2020. Mitochondrial pathways and respiratory control: An Introduction to OXPHOS Analysis. 5th ed. Bioenergetics Communications 2020, 2–2. 10.26124/bec:2020-0002

13. Gutierrez, E., Wiggins, D., Fielding, B., Gould, A.P., 2007. Specialized hepatocyte-like cells regulate Drosophila lipid metabolism. Nature 445, 275–280. 10.1038/nature05382

14. Hafen, P.S., Preece, C.N., Sorensen, J.R., Hancock, C.R., Hyldahl, R.D., 2018. Repeated exposure to heat stress induces mitochondrial adaptation in human skeletal muscle. J Appl Physiol (1985) 125, 1447–1455. 10.1152/japplphysiol.00383.2018

15. Humez, S., Monet, M., van Coppenolle, F., Delcourt, P., Prevarskaya, N., 2004. The role of intracellular pH in cell growth arrest induced by ATP. Am J Physiol Cell Physiol 287, C1733–1746. 10.1152/ajpcell.00578.2003

16. Jalal, M., Andersen, T., Hessen, D.O., 2015. Temperature and developmental responses of body and cell size in Drosophila; effects of polyploidy and genome configuration. J Therm Biol 51, 1–14. 10.1016/j.jtherbio.2015.02.011

17. Jandova, J., Shi, M., Norman, K.G., Stricklin, G.P., Sligh, J.E., 2012. Somatic alterations in mitochondrial DNA produce changes in cell growth and metabolism supporting a tumorigenic phenotype. Biochimica et Biophysica Acta (BBA) - Molecular Basis of Disease 1822, 293–300. 10.1016/j.bbadis.2011.11.010

18. Jevtov, I., Zacharogianni, M., van Oorschot, M.M., van Zadelhoff, G., Aguilera-Gomez, A., Vuillez, I., Braakman, I., Hafen, E., Stocker, H., Rabouille, C., 2015. TORC2 mediates the heat stress response in Drosophila by promoting the formation of stress granules. J Cell Sci 128, 2497–2508. 10.1242/jcs.168724

19. Klepsatel, P., Wildridge, D., Gáliková, M., 2019. Temperature induces changes in Drosophila energy stores. Sci Rep 9, 5239. 10.1038/s41598-019-41754-5

20. Liu, C.-T., Brooks, G.A., 2011. Mild heat stress induces mitochondrial biogenesis in C2C12 myotubes. Journal of Applied Physiology 112, 354. 10.1152/japplphysiol.00989.2011

21. Lu, J., Li, H., Yu, D., Zhao, P., Liu, Y., 2023a. Heat stress inhibits the proliferation and differentiation of myoblasts and is associated with damage to mitochondria. Front. Cell Dev. Biol. 11. 10.3389/fcell.2023.1171506

22. Lu, J., Li, H., Yu, D., Zhao, P., Liu, Y., 2023b. Heat stress inhibits the proliferation and differentiation of myoblasts and is associated with damage to mitochondria. Front. Cell Dev. Biol. 11. 10.3389/fcell.2023.1171506

23. Maunder, E., King, A., Rothschild, J.A., Brick, M.J., Leigh, W.B., Hedges, C.P., Merry, T.L., Kilding, A.E., 2024. Locally applied heat stress during exercise training may promote adaptations to mitochondrial enzyme activities in skeletal muscle. Pflugers Arch - Eur J Physiol 476, 939–948. 10.1007/s00424-024-02939-8

24. Miettinen, T.P., Björklund, M., 2017. Mitochondrial Function and Cell Size: An Allometric Relationship. Trends Cell Biol 27, 393–402. 10.1016/j.tcb.2017.02.006

25. Miettinen, T.P., Caldez, M.J., Kaldis, P., Björklund, M., 2017. Cell size control – a mechanism for maintaining fitness and function. BioEssays 39, 1700058. 10.1002/bies.201700058

26. Mishra, P., Pandey, C.M., Singh, U., Gupta, A., Sahu, C., Keshri, A., 2019. Descriptive Statistics and Normality Tests for Statistical Data. Ann Card Anaesth 22, 67–72. 10.4103/aca.ACA_157_18

27. Moraes, A.M., Jorge, S.A.C., Astray, R.M., Suazo, C.A.T., Calderón Riquelme, C.E., Augusto, E.F.P., Tonso, A., Pamboukian, M.M., Piccoli, R.A.M., Barral, M.F., Pereira, C.A., 2012. Drosophila melanogaster S2 cells for expression of heterologous genes: From gene cloning to bioprocess development. Biotechnol Adv 30, 613–628. 10.1016/j.biotechadv.2011.10.009

28. Nadeau, E.A.W., Teets, N.M., 2020. Evidence for a rapid cold hardening response in cultured Drosophila S2 cells. J Exp Biol 223, jeb212613. 10.1242/jeb.212613

29. Nicholls, D.G., 2009. Spare respiratory capacity, oxidative stress and excitotoxicity. Biochem Soc Trans 37, 1385–1388. 10.1042/BST0371385

30. Pamboukian, M.M., Jorge, S.A.C., Santos, M.G., Yokomizo, A.Y., Pereira, C.A., Tonso, A., 2008. Insect cells respiratory activity in bioreactor. Cytotechnology 57, 37–44. 10.1007/s10616-007-9118-8

31. Pandey, U.B., Nichols, C.D., 2011. Human disease models in Drosophila melanogaster and the role of the fly in therapeutic drug discovery. Pharmacol Rev 63, 411–436. 10.1124/pr.110.003293

32. Pesta, D., Gnaiger, E., 2012. High-resolution respirometry: OXPHOS protocols for human cells and permeabilized fibers from small biopsies of human muscle. Methods Mol Biol 810, 25–58. 10.1007/978-1-61779-382-0_3

33. Place, T.L., Domann, F.E., Case, A.J., 2017. Limitations of oxygen delivery to cells in culture: An underappreciated problem in basic and translational research. Free Radic Biol Med 113, 311–322. 10.1016/j.freeradbiomed.2017.10.003

34. Rao, P.N., Engelberg, J., 1965. HeLa Cells: Effects of Temperature on the Life Cycle. Science 148, 1092–1094. 10.1126/science.148.3673.1092

35. Roussel, D., Voituron, Y., 2020. Mitochondrial Costs of Being Hot: Effects of Acute Thermal Change on Liver Bioenergetics in Toads (Bufo bufo). Front. Physiol. 11. 10.3389/fphys.2020.00153

36. Schneider, I., 1972. Cell lines derived from late embryonic stages of Drosophila melanogaster. J Embryol Exp Morphol 27, 353–365.

37. Schneider’s Drosophila Line 2 [D. Mel. (2), SL2] - CRL-1963 | ATCC [WWW Document], n.d. . https://www.atcc.org. URL https://www.atcc.org/products/crl-1963 (accessed 11.6.25).

38. Seel, A., Padovani, F., Mayer, M., Finster, A., Bureik, D., Thoma, F., Osman, C., Klecker, T., Schmoller, K.M., 2023. Regulation with cell size ensures mitochondrial DNA homeostasis during cell growth. Nat Struct Mol Biol 30, 1549–1560. 10.1038/s41594-023-01091-8

39. Shao-Hua, C., Hong-Liang, S., Zuo-Hu, L., 1998. Effect of Temperature Oscillation on Insect Cell Growth and Baculovirus Replication. Applied and Environmental Microbiology 64, 2237–2239. 10.1128/AEM.64.6.2237-2239.1998

40. Sokolova, I.M., 2023. Ectotherm mitochondrial economy and responses to global warming. Acta Physiol (Oxf) 237, e13950. 10.1111/apha.13950

41. statsmodels n.d. Tukey HSD posthoc test for group interactions in two-way ANOVA · Issue #6799 · statsmodels/statsmodels [WWW Document]. GitHub. URL https://github.com/statsmodels/statsmodels/issues/6799 (accessed 11.6.25).

42. Terzioglu, M., Veeroja, K., Montonen, T., Ihalainen, T.O., Salminen, T.S., Bénit, P., Rustin, P., Chang, Y.-T., Nagai, T., Jacobs, H.T., 2023. Mitochondrial temperature homeostasis resists external metabolic stresses. eLife 12. 10.7554/eLife.89232.2

43. Thoral, E., Correia, M.G., Chamkha, I., Elmér, E., Nord, A., 2025. Incubation temperature shapes growth and mitochondrial metabolism across embryonic development in Japanese quail. Proc Biol Sci 292, 20251752. 10.1098/rspb.2025.1752

44. Vander Heiden, M.G., Plas, D.R., Rathmell, J.C., Fox, C.J., Harris, M.H., Thompson, C.B., 2001. Growth factors can influence cell growth and survival through effects on glucose metabolism. Mol Cell Biol 21, 5899–5912. 10.1128/MCB.21.17.5899-5912.2001

45. Von Schulze, A.T., Geiger, P.C., 2022. Heat and mitochondrial bioenergetics. Current Opinion in Physiology 27, 100553. 10.1016/j.cophys.2022.100553

46. Yadava, N., Nicholls, D.G., 2007. Spare Respiratory Capacity Rather Than Oxidative Stress Regulates Glutamate Excitotoxicity after Partial Respiratory Inhibition of Mitochondrial Complex I with Rotenone. J Neurosci 27, 7310–7317. 10.1523/JNEUROSCI.0212-07.2007

47. Yao, C.-H., Wang, R., Wang, Y., Kung, C.-P., Weber, J.D., Patti, G.J., 2019. Mitochondrial fusion supports increased oxidative phosphorylation during cell proliferation. eLife 8, e41351. 10.7554/eLife.41351

