## Supplementary tables S1-S14 for "Effects of cell growth and temperature on mitochondrial physiology of *Drosophila melanogaster* embryonic cell line"

### Slide 1
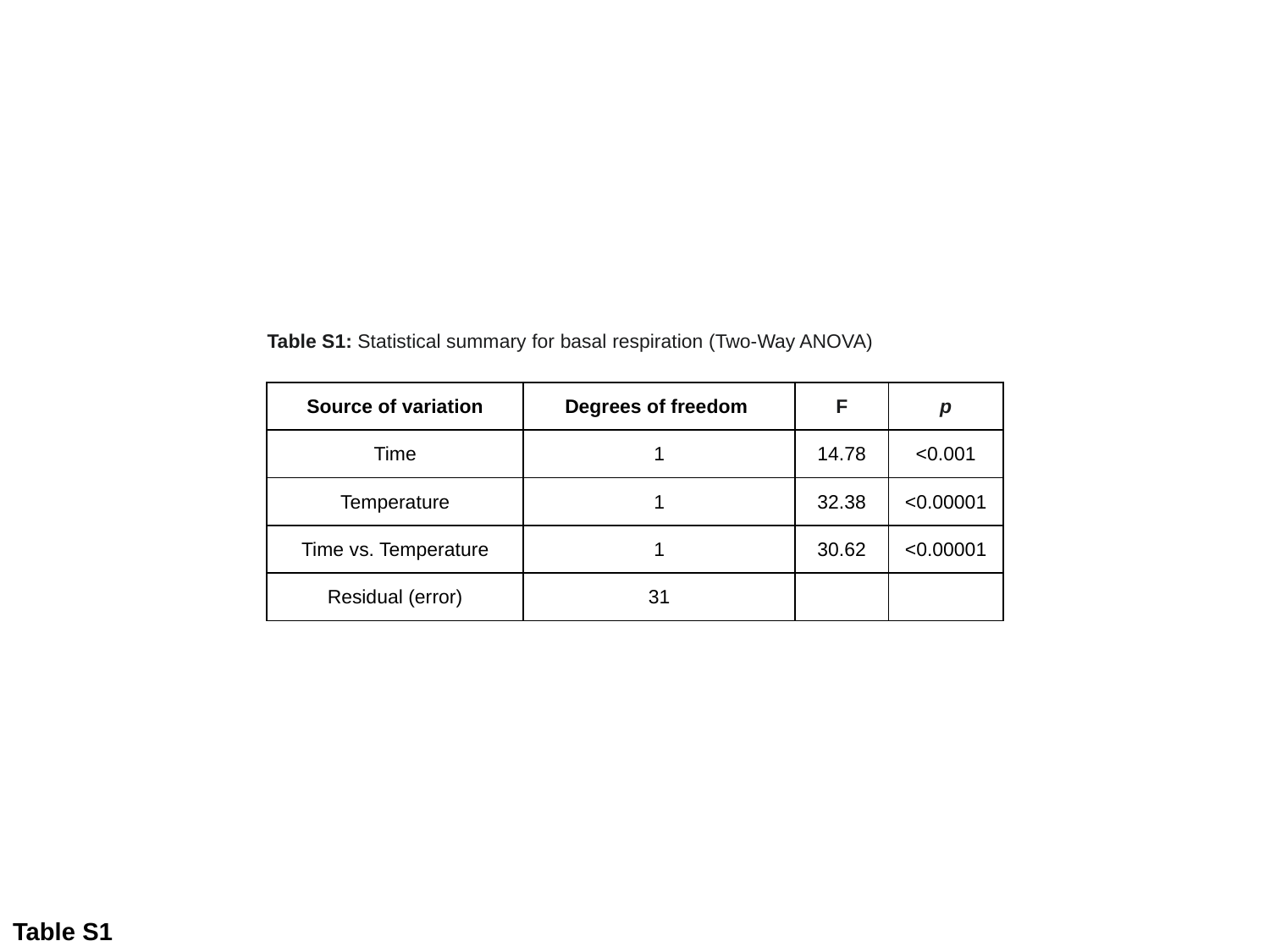

Table S1: Statistical summary for basal respiration (Two-Way ANOVA)
| Source of variation | Degrees of freedom | F | p |
| --- | --- | --- | --- |
| Time | 1 | 14.78 | <0.001 |
| Temperature | 1 | 32.38 | <0.00001 |
| Time vs. Temperature | 1 | 30.62 | <0.00001 |
| Residual (error) | 31 | | |
Table S1

### Slide 2
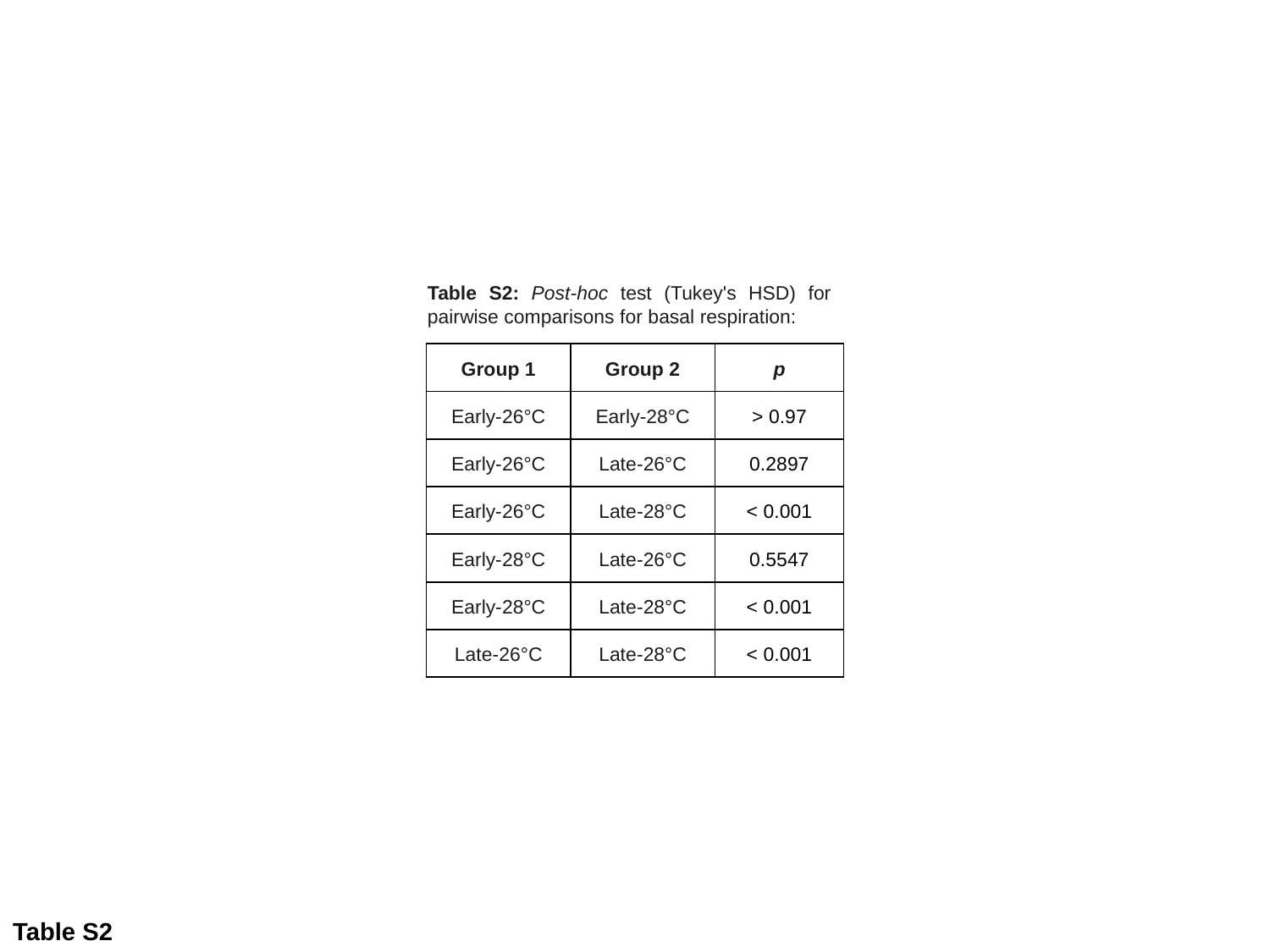

Table S2: Post-hoc test (Tukey's HSD) for pairwise comparisons for basal respiration:
| Group 1 | Group 2 | p |
| --- | --- | --- |
| Early-26°C | Early-28°C | > 0.97 |
| Early-26°C | Late-26°C | 0.2897 |
| Early-26°C | Late-28°C | < 0.001 |
| Early-28°C | Late-26°C | 0.5547 |
| Early-28°C | Late-28°C | < 0.001 |
| Late-26°C | Late-28°C | < 0.001 |
Table S2

### Slide 3
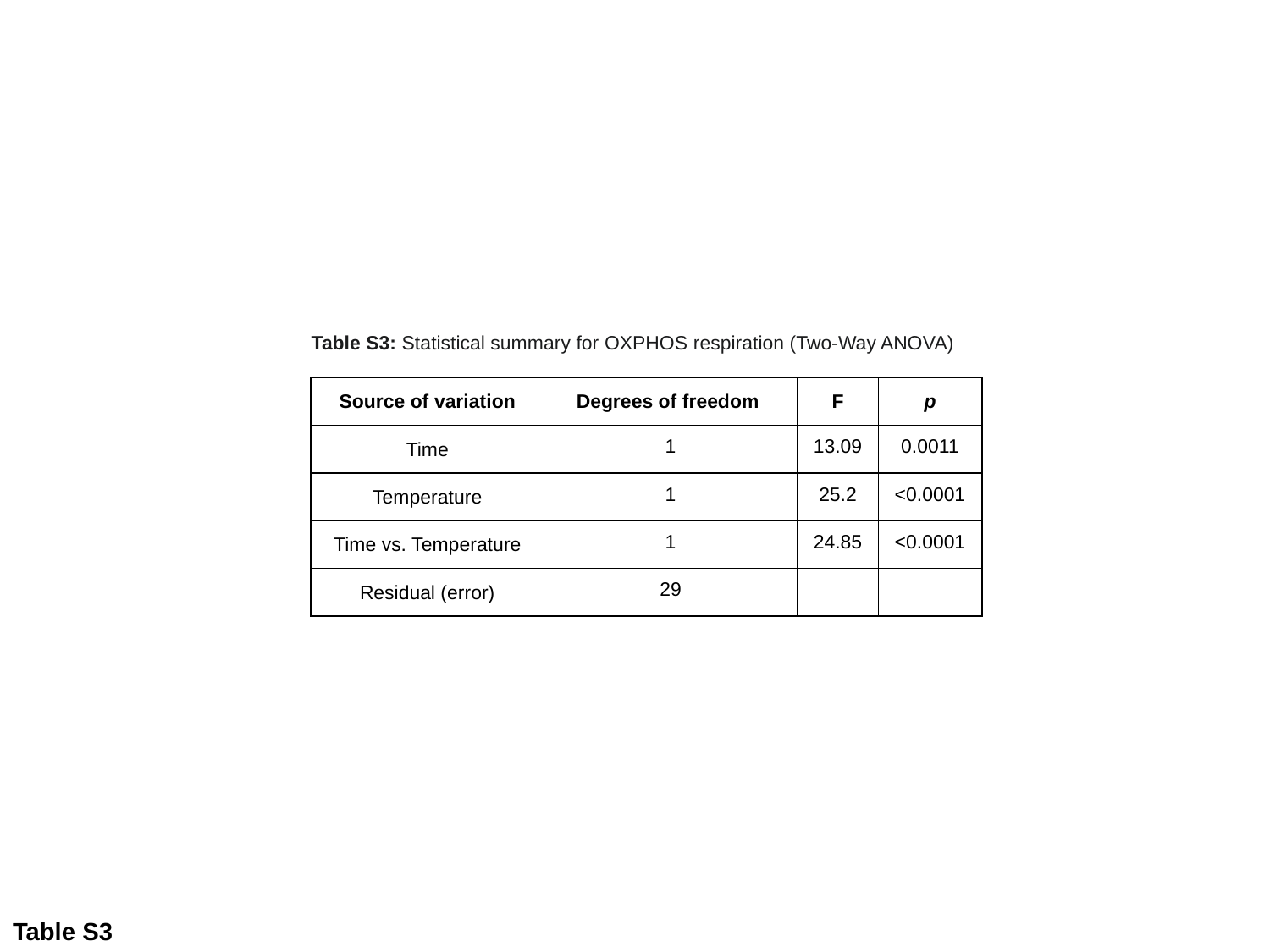

Table S3: Statistical summary for OXPHOS respiration (Two-Way ANOVA)
| Source of variation | Degrees of freedom | F | p |
| --- | --- | --- | --- |
| Time | 1 | 13.09 | 0.0011 |
| Temperature | 1 | 25.2 | <0.0001 |
| Time vs. Temperature | 1 | 24.85 | <0.0001 |
| Residual (error) | 29 | | |
Table S3

### Slide 4
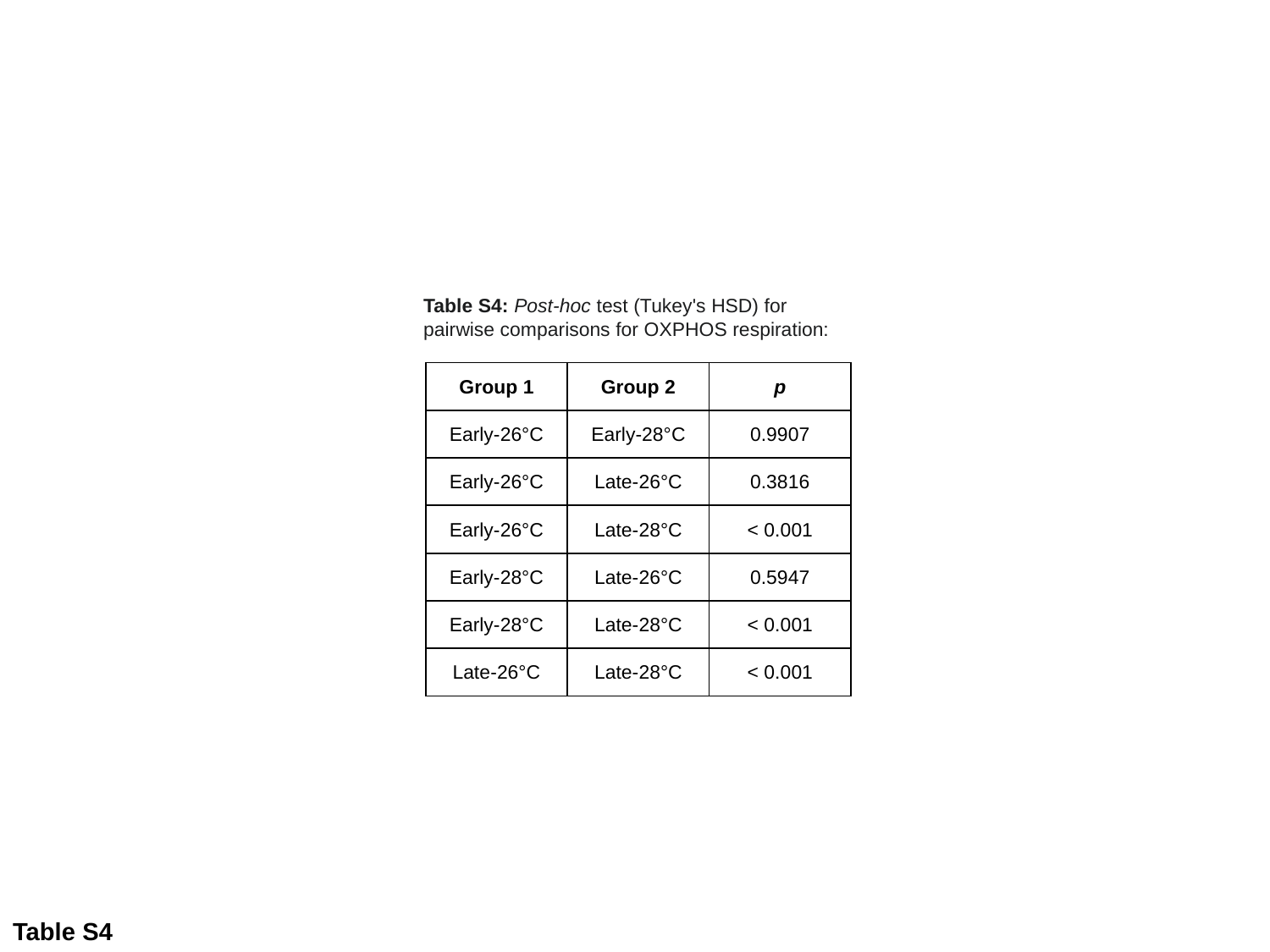

Table S4: Post-hoc test (Tukey's HSD) for pairwise comparisons for OXPHOS respiration:
| Group 1 | Group 2 | p |
| --- | --- | --- |
| Early-26°C | Early-28°C | 0.9907 |
| Early-26°C | Late-26°C | 0.3816 |
| Early-26°C | Late-28°C | < 0.001 |
| Early-28°C | Late-26°C | 0.5947 |
| Early-28°C | Late-28°C | < 0.001 |
| Late-26°C | Late-28°C | < 0.001 |
Table S4

### Slide 5
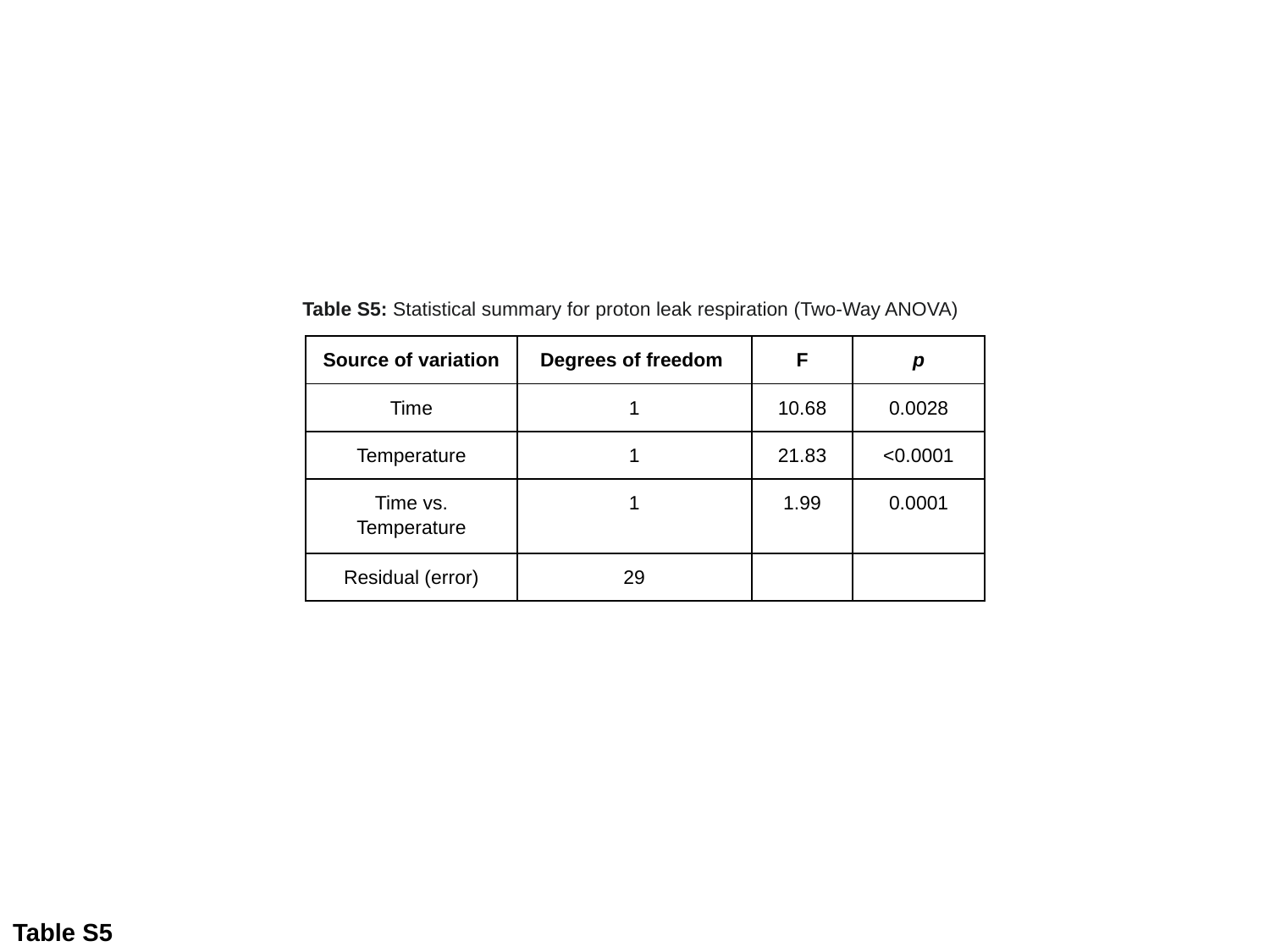

Table S5: Statistical summary for proton leak respiration (Two-Way ANOVA)
| Source of variation | Degrees of freedom | F | p |
| --- | --- | --- | --- |
| Time | 1 | 10.68 | 0.0028 |
| Temperature | 1 | 21.83 | <0.0001 |
| Time vs. Temperature | 1 | 1.99 | 0.0001 |
| Residual (error) | 29 | | |
Table S5

### Slide 6
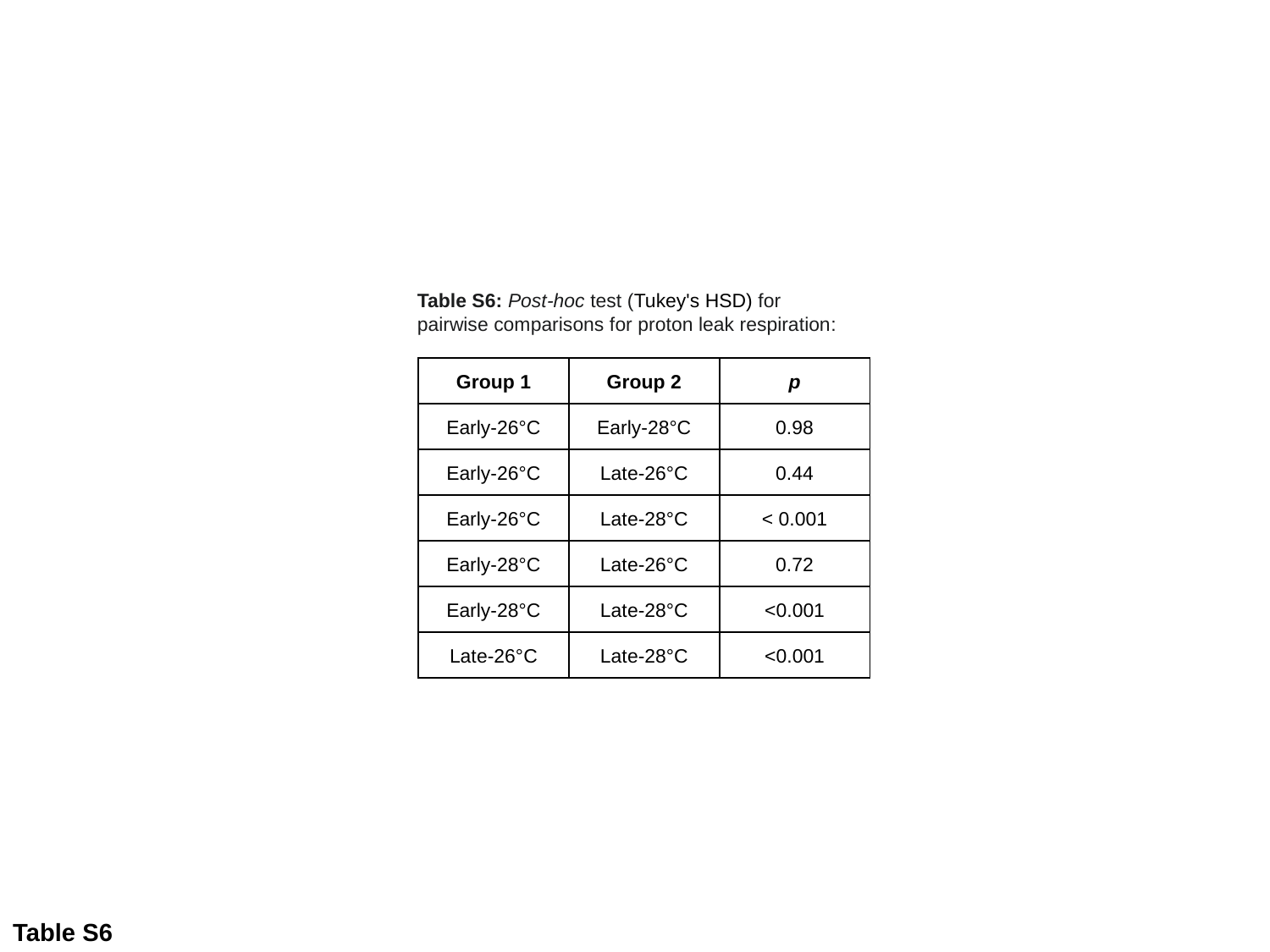

Table S6: Post-hoc test (Tukey's HSD) for pairwise comparisons for proton leak respiration:
| Group 1 | Group 2 | p |
| --- | --- | --- |
| Early-26°C | Early-28°C | 0.98 |
| Early-26°C | Late-26°C | 0.44 |
| Early-26°C | Late-28°C | < 0.001 |
| Early-28°C | Late-26°C | 0.72 |
| Early-28°C | Late-28°C | <0.001 |
| Late-26°C | Late-28°C | <0.001 |
Table S6

### Slide 7
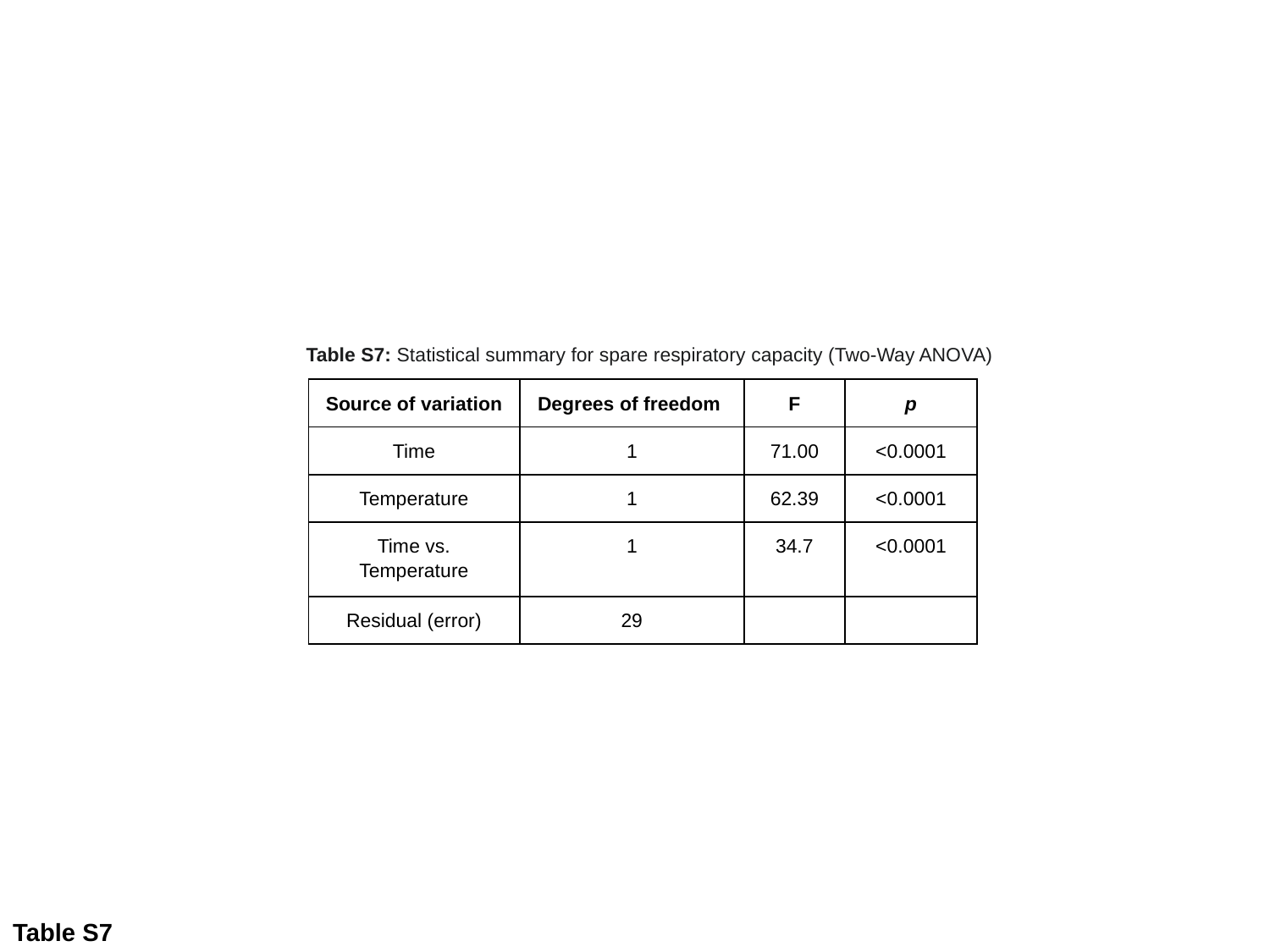

Table S7: Statistical summary for spare respiratory capacity (Two-Way ANOVA)
| Source of variation | Degrees of freedom | F | p |
| --- | --- | --- | --- |
| Time | 1 | 71.00 | <0.0001 |
| Temperature | 1 | 62.39 | <0.0001 |
| Time vs. Temperature | 1 | 34.7 | <0.0001 |
| Residual (error) | 29 | | |
Table S7

### Slide 8
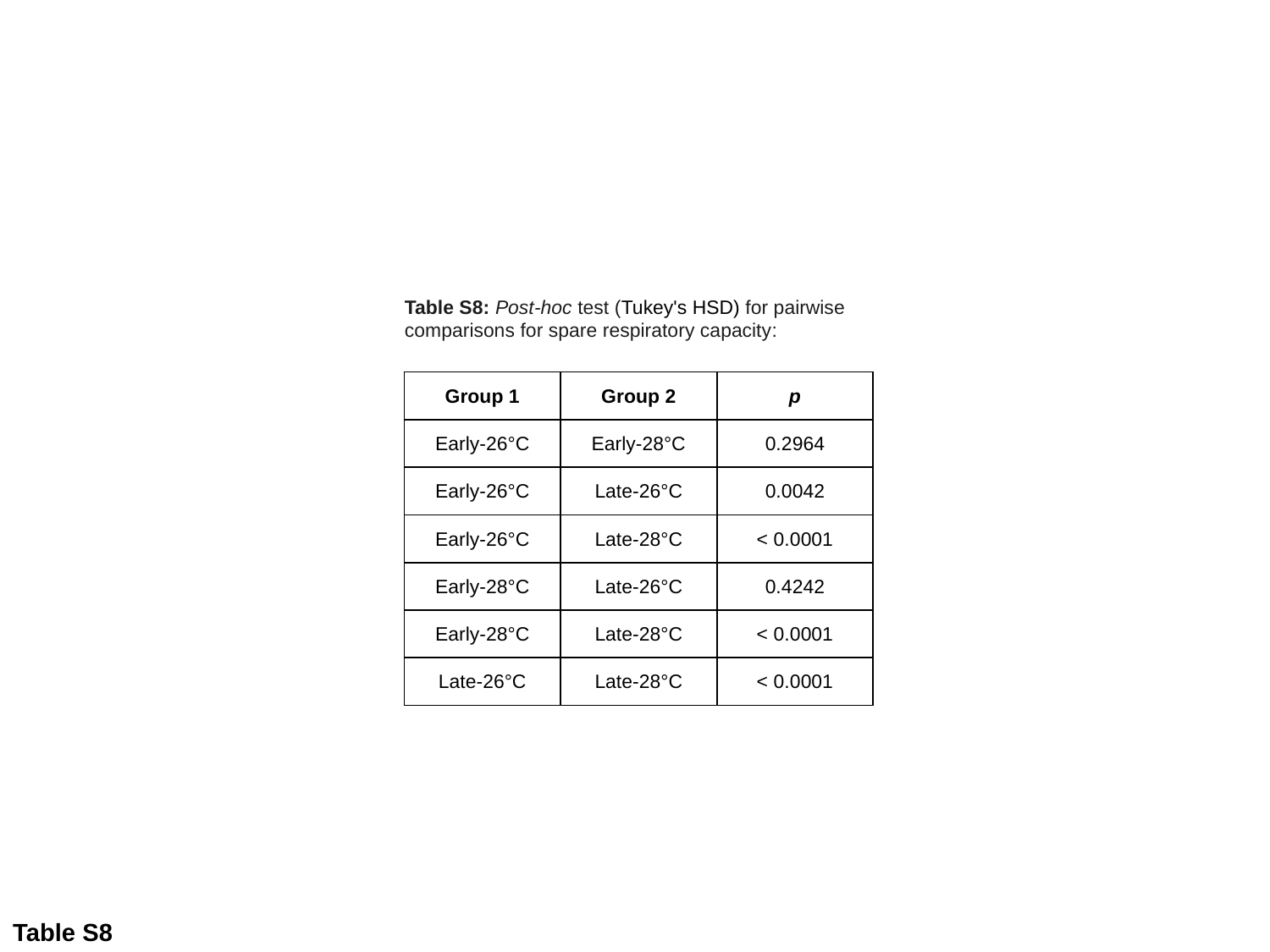

Table S8: Post-hoc test (Tukey's HSD) for pairwise comparisons for spare respiratory capacity:
| Group 1 | Group 2 | p |
| --- | --- | --- |
| Early-26°C | Early-28°C | 0.2964 |
| Early-26°C | Late-26°C | 0.0042 |
| Early-26°C | Late-28°C | < 0.0001 |
| Early-28°C | Late-26°C | 0.4242 |
| Early-28°C | Late-28°C | < 0.0001 |
| Late-26°C | Late-28°C | < 0.0001 |
Table S8

### Slide 9
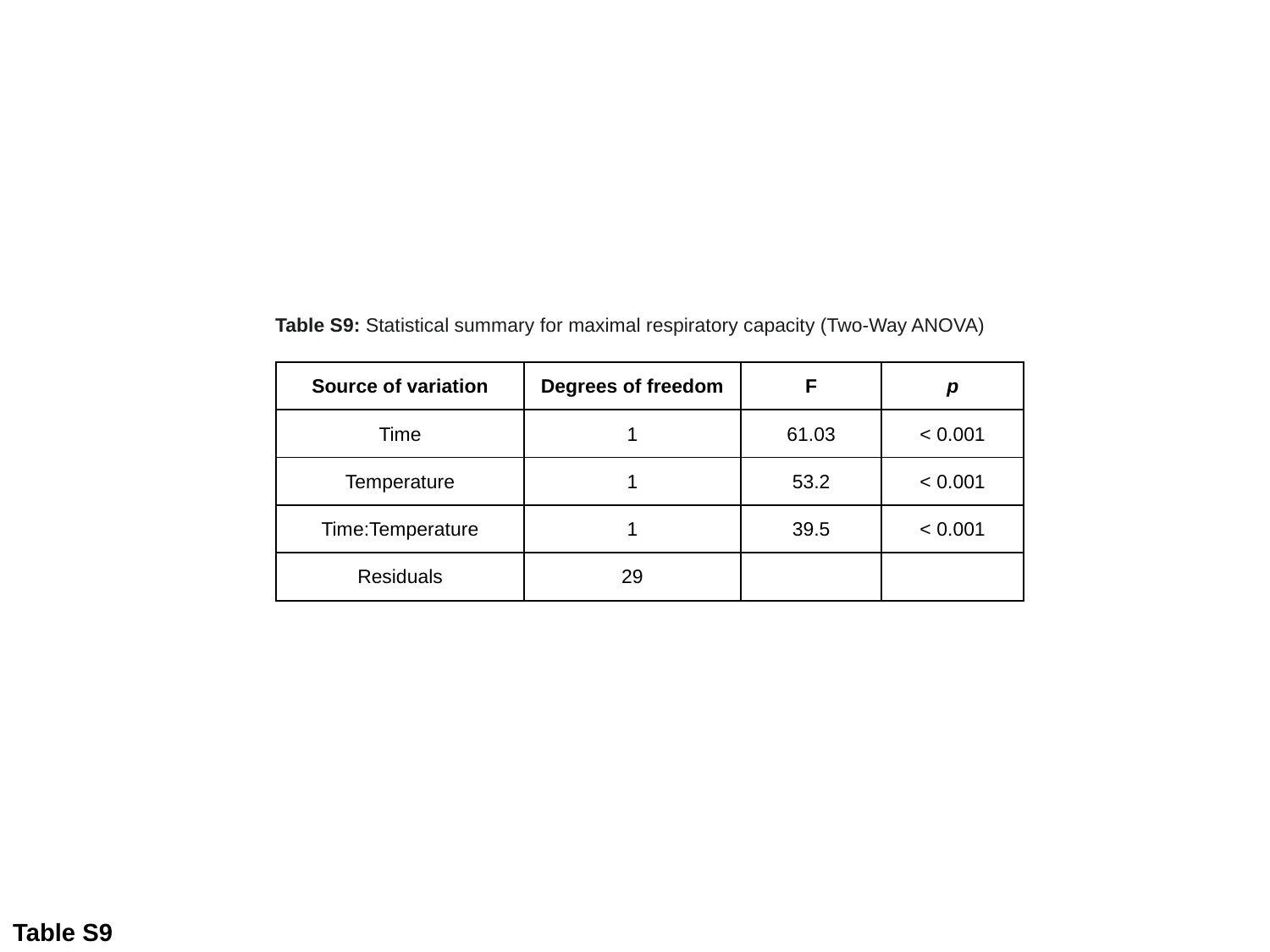

Table S9: Statistical summary for maximal respiratory capacity (Two-Way ANOVA)
| Source of variation | Degrees of freedom | F | p |
| --- | --- | --- | --- |
| Time | 1 | 61.03 | < 0.001 |
| Temperature | 1 | 53.2 | < 0.001 |
| Time:Temperature | 1 | 39.5 | < 0.001 |
| Residuals | 29 | | |
Table S9

### Slide 10
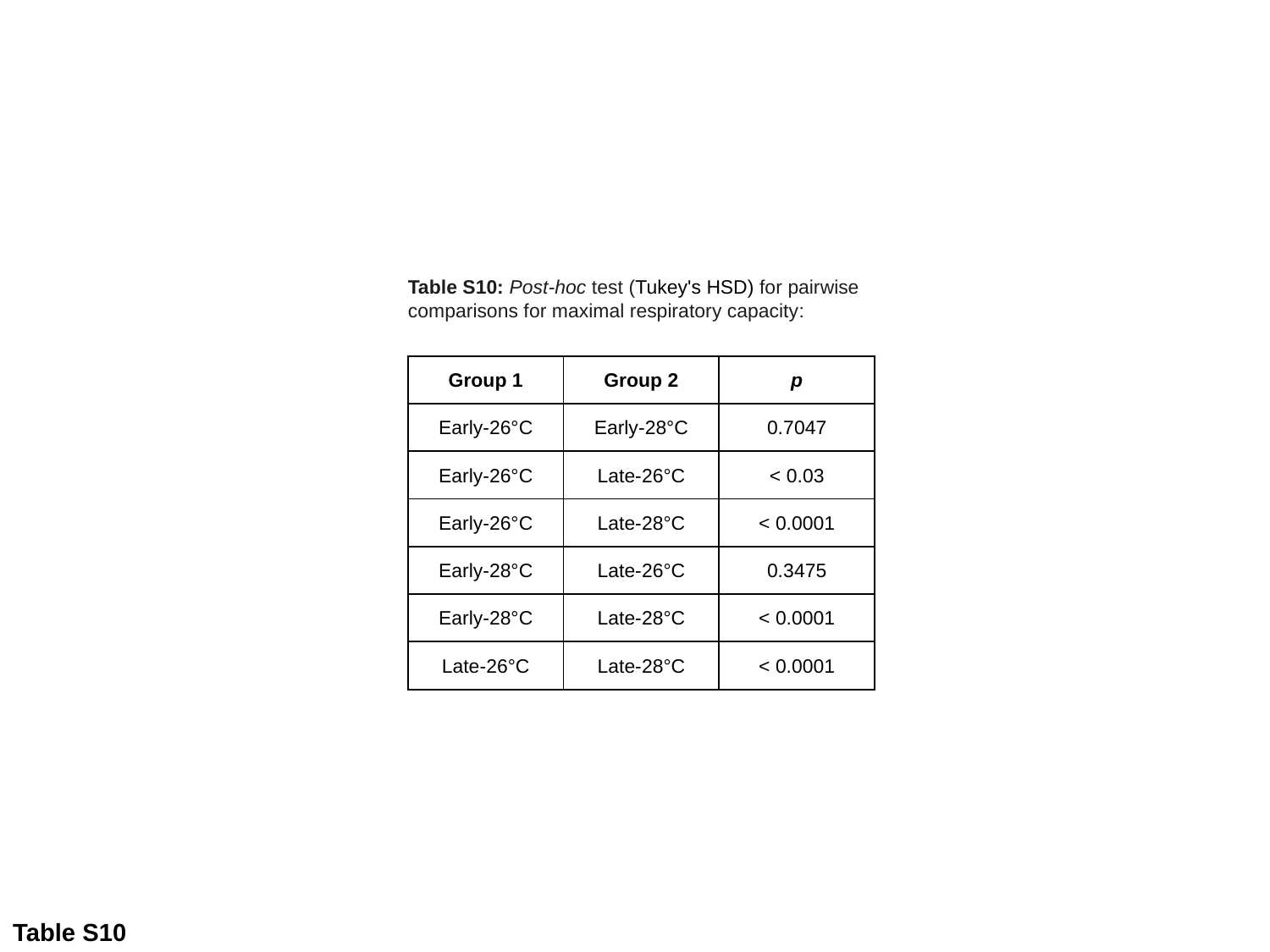

Table S10: Post-hoc test (Tukey's HSD) for pairwise comparisons for maximal respiratory capacity:
| Group 1 | Group 2 | p |
| --- | --- | --- |
| Early-26°C | Early-28°C | 0.7047 |
| Early-26°C | Late-26°C | < 0.03 |
| Early-26°C | Late-28°C | < 0.0001 |
| Early-28°C | Late-26°C | 0.3475 |
| Early-28°C | Late-28°C | < 0.0001 |
| Late-26°C | Late-28°C | < 0.0001 |
Table S10

### Slide 11
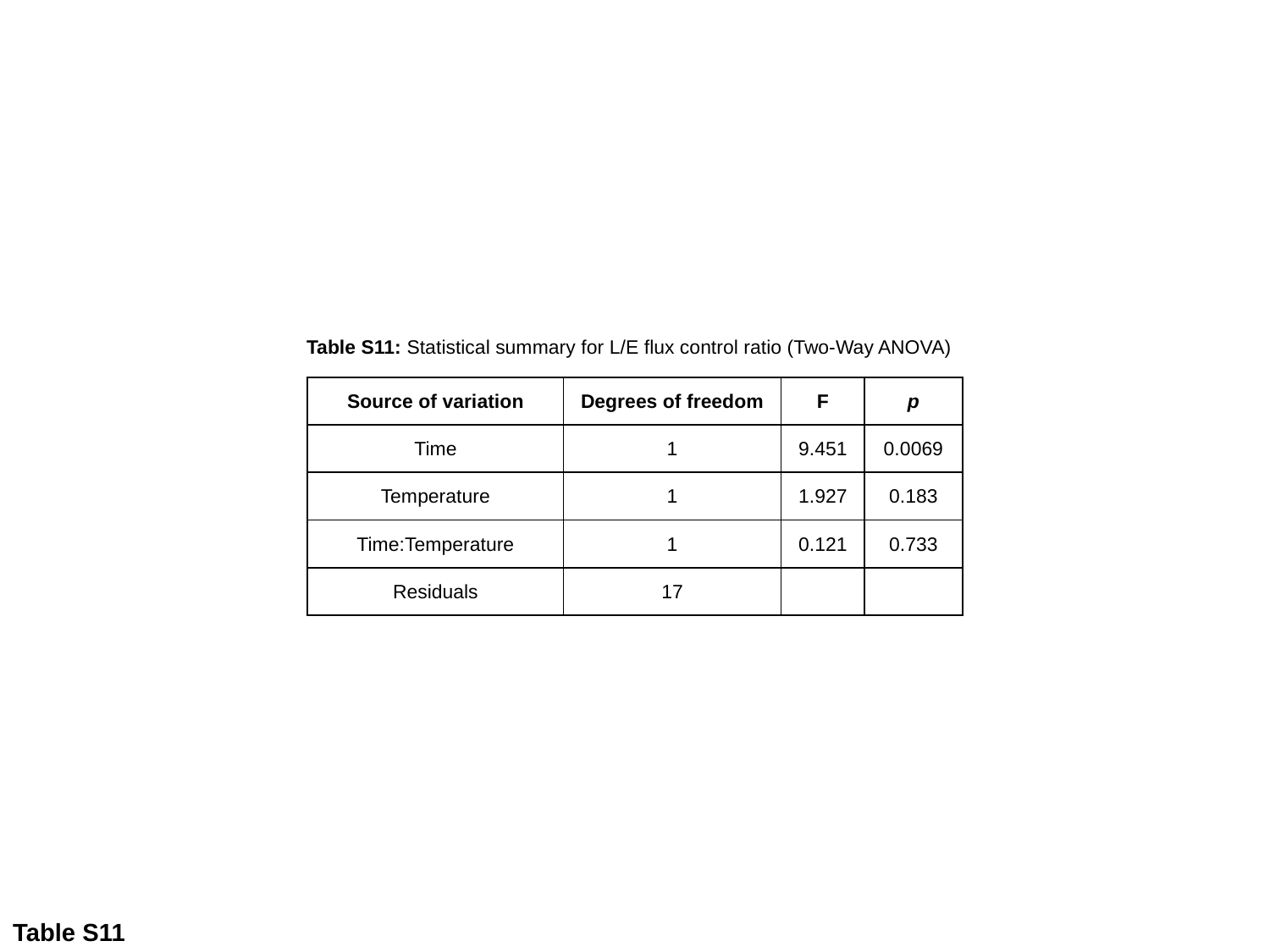

Table S11: Statistical summary for L/E flux control ratio (Two-Way ANOVA)
| Source of variation | Degrees of freedom | F | p |
| --- | --- | --- | --- |
| Time | 1 | 9.451 | 0.0069 |
| Temperature | 1 | 1.927 | 0.183 |
| Time:Temperature | 1 | 0.121 | 0.733 |
| Residuals | 17 | | |
Table S11

### Slide 12
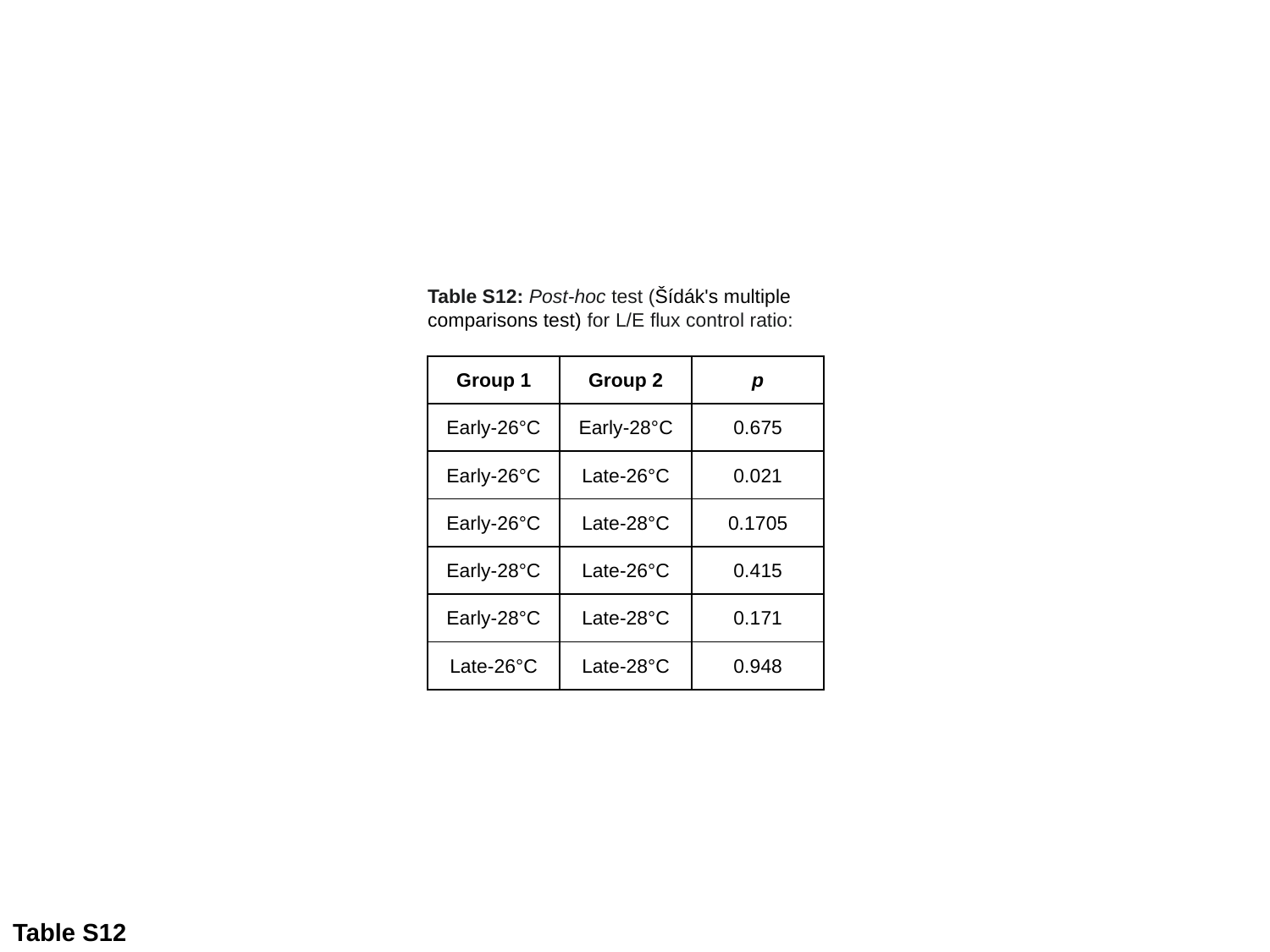

Table S12: Post-hoc test (Šídák's multiple comparisons test) for L/E flux control ratio:
| Group 1 | Group 2 | p |
| --- | --- | --- |
| Early-26°C | Early-28°C | 0.675 |
| Early-26°C | Late-26°C | 0.021 |
| Early-26°C | Late-28°C | 0.1705 |
| Early-28°C | Late-26°C | 0.415 |
| Early-28°C | Late-28°C | 0.171 |
| Late-26°C | Late-28°C | 0.948 |
Table S12

### Slide 13
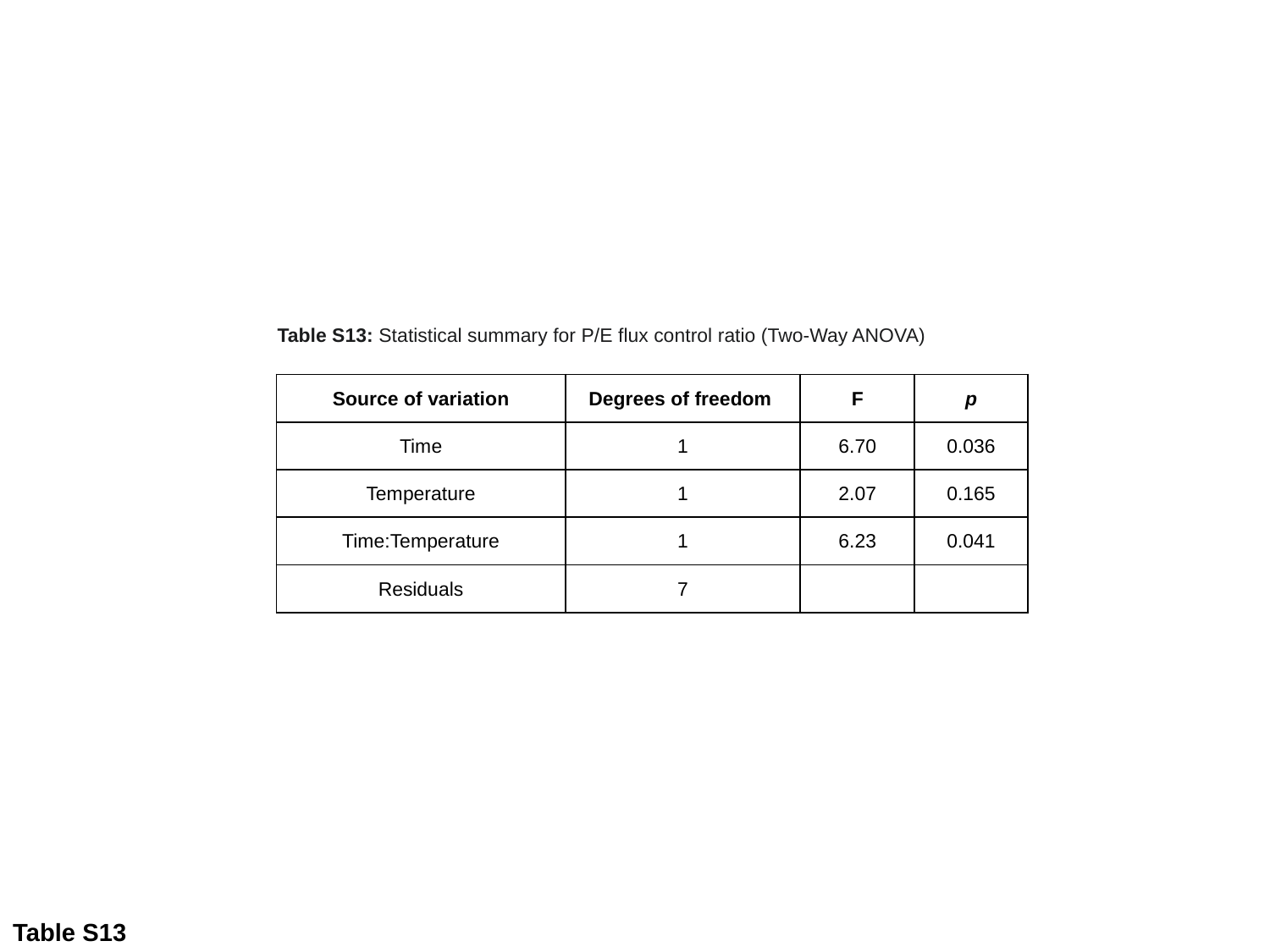

Table S13: Statistical summary for P/E flux control ratio (Two-Way ANOVA)
| Source of variation | Degrees of freedom | F | p |
| --- | --- | --- | --- |
| Time | 1 | 6.70 | 0.036 |
| Temperature | 1 | 2.07 | 0.165 |
| Time:Temperature | 1 | 6.23 | 0.041 |
| Residuals | 7 | | |
Table S13

### Slide 14
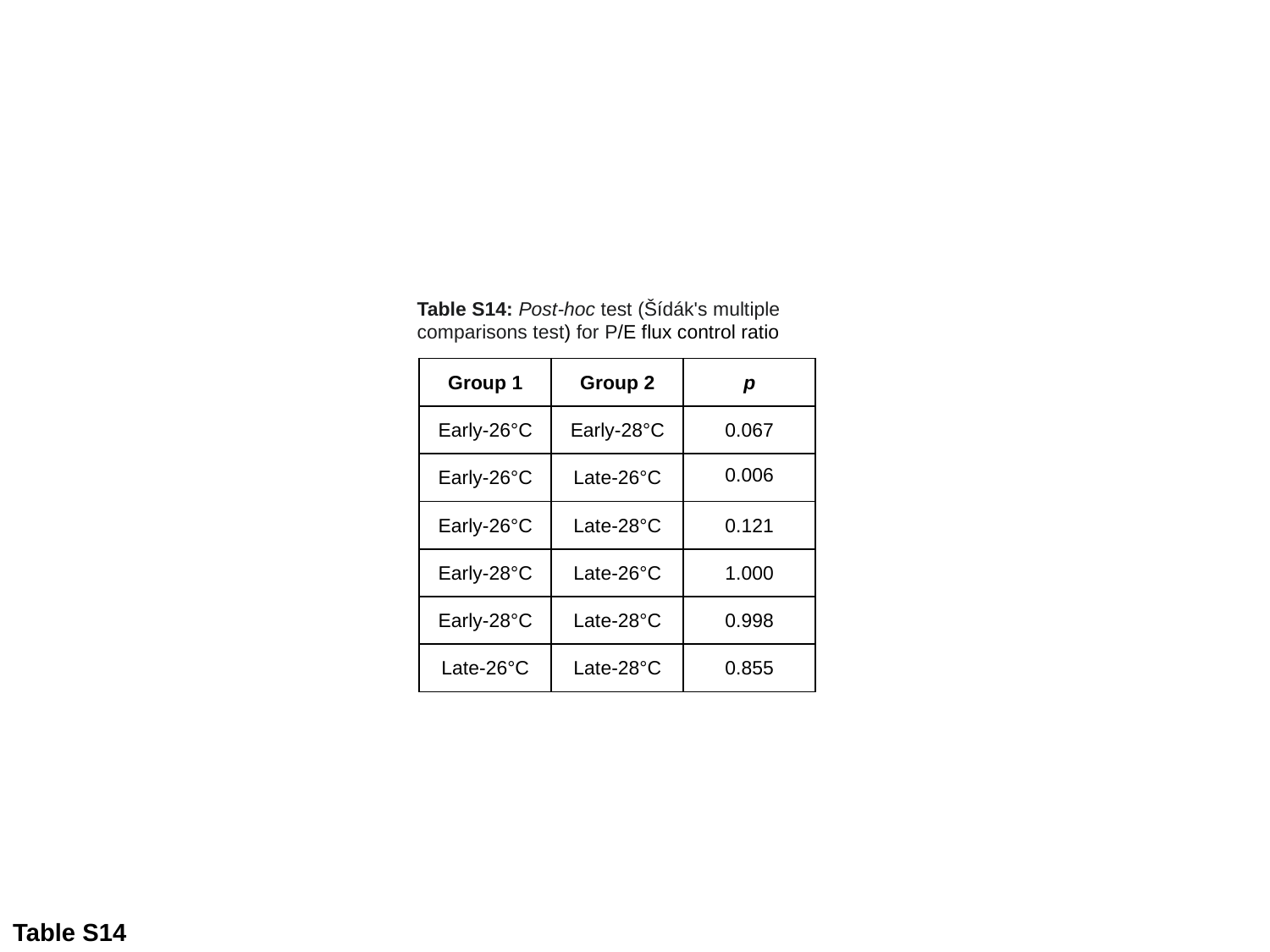

Table S14: Post-hoc test (Šídák's multiple comparisons test) for P/E flux control ratio
| Group 1 | Group 2 | p |
| --- | --- | --- |
| Early-26°C | Early-28°C | 0.067 |
| Early-26°C | Late-26°C | 0.006 |
| Early-26°C | Late-28°C | 0.121 |
| Early-28°C | Late-26°C | 1.000 |
| Early-28°C | Late-28°C | 0.998 |
| Late-26°C | Late-28°C | 0.855 |
Table S14
