## Supplementary material for "Effects of cell growth and temperature on mitochondrial physiology of *Drosophila melanogaster* embryonic cell line": AI prompt for statistical analyses

**Prompt designed for multiple comparison analyses:**

Objective: Perform a comprehensive statistical analysis of a 2x2 factorial dataset from the attached file (data.xls) and generate a publishable-quality report.

**1. Data and Variables:**

Attached File: data.xls

Independent Variables (Factors):

Time: 2 levels (Early, Late)

Temperature: 2 levels (26°C, 28°C)

**Experimental Groups (4 total):**

Early / 26°C

Early / 28°C

Late / 26°C

Late / 28°C

**Dependent Variables (Measures):**

**Group 1 (Respirometry):**

Basal

OXPHOS (ATP-linked respiration)

Proton Leak

Spare Respiratory Capacity

Maximal Electron Transfer System (ETS)

**Group 2 (Bioenergetic Efficiency Ratios):**

L/E (Leak/Maximal)

P/E (ATP-linked/Maximal)

**2. Data Parsing Instructions:**

Parse the data.xls file. The data is organized into columns corresponding to the 7 dependent variables and the 4 experimental groups.

The user notes on color-coding (red/blue/green/black fonts) and shaded boxes are human-facing metadata. You must identify the data by matching the column headers for the dependent variables and the group labels (Time and Temperature).

All analyses must be performed separately for Group 1 (Respirometry) and Group 2 (Bioenergetic Efficiency Ratios). Do not statistically compare variables between these two groups (e.g., do not compare 'Basal' to 'L/E').

**3. Statistical Analysis Workflow:**

For each of the 7 dependent variables (Basal, OXPHOS, ..., L/E, P/E), perform the following sequence:

Assumption Testing:

Test the data for each of the 4 experimental groups for normality (e.g., Shapiro-Wilk test).

Test for homogeneity of variances across the groups (e.g., Levene's test).

Primary Statistical Test (Conditional):

Parametric Path: If the data meets the assumptions of normality and homogeneity of variance, perform a Two-Way ANOVA + tukey´s post hoc tests.

Non-Parametric Path: If the data violate these assumptions, perform a robust non-parametric test suitable for a two-way factorial design that can test for interactions (e.g., kruskal-wallis + dunn´s post hoc tests).

Reported Effects: For the chosen test (either Two-Way ANOVA or kruskal-wallis), you must determine and report the statistics (F-statistic, p-value, etc.) for:

Main Effect of Time

Main Effect of Temperature

Time x Temperature Interaction Effect

Post-Hoc Analysis:

If a significant main effect or interaction is found, perform a post-hoc test to compare the means (or medians) of all four experimental groups.

If Two-Way ANOVA was used: Use Tukey's HSD test.

If a non-parametric test was used: Use an appropriate corresponding post-hoc test (e.g., Dunn's test or pairwise comparisons on the transformed data, with p-values adjusted for multiple comparisons like Bonferroni).

**4. Required Deliverables:**

**A. Report Document (.docx file)**

Structure the document with clear sections for each of the 7 dependent variables.

For each variable, provide:

Results (Text): A written summary of the findings. State the results of the assumption tests, which primary test was used, and interpret the main effects and interaction (e.g., "A Two-Way ANOVA revealed a significant main effect for Time (F(df1, df2) = X, p = Y) and a significant Time x Temperature interaction (F(df1, df2) = Z, p = Q)...").

Summary Table (Publishable): A formatted table (e.g., an ANOVA table) summarizing the results of the primary test. Include columns for Source of Variation (Time, Temperature, Time x Temperature, Error/Residual), Degrees of Freedom, F-statistic (or equivalent), and p-value.

Post-Hoc Table (Publishable): A table clearly presenting the pairwise comparisons from the post-hoc test, showing which groups are significantly different from one another.
